# Disruption of glycogen metabolism alters cell size in *Escherichia coli*

**DOI:** 10.1101/2020.06.01.127233

**Authors:** Felix E van der Walt, Léo Bürgy, Lize Engelbrecht, Rozanne CM Adams, Jessica de Stadler, Lindi Strydom, Gavin M George, Samuel C Zeeman, Jens Kossmann, James R Lloyd

## Abstract

The availability of nutrients impacts cell size and growth rate in many organisms. Research in *E. coli* has traditionally focused on the influence of exogenous nutrient sources on cell size through their effect on growth and cell cycle progression. Utilising a set of mutants where three genes involved in glycogen degradation - glycogen phosphorylase (*glgP)*, glycogen debranching enzyme (*glgX*) and maltodextrin phosphorylase (*malP)* - were disrupted, we examined if endogenous polyglucan degradation affects cell size. It was found that mutations to *malP* increased cell lengths and resulted in substantial heterogeneity of cell size. This was most apparent during exponential growth and the phenotype was unaccompanied by alterations in Z-ring occurrence, cellular FtsZ levels and generation times. Δ*malP* mutant cells did, however, accumulate increased DnaA amounts at late growth stages indicating a potential effect on DNA replication. Replication run-out experiments demonstrated that this was indeed the case, and that DNA replication was also affected in the other mutants. Bacteria with a disruption in *glgX* accumulated glycogen and protein inclusion bodies that coincided with each other at inter-nucleoid and polar regions.

## INTRODUCTION

Robust adaptive survival strategies are employed by bacteria to promote fitness. Most notably, when nutrients are abundant, bacterial cells demonstrate increases in length, width and growth rate^1^ which permit parental cells to promote vigour of future generations by giving rise to larger daughter cells^2^. Enhanced rates of macromolecular biosynthesis accompany nutrient-dependent size increases to ensure that larger cells are born with more DNA, RNA and protein^3–6^. This positive scaling relationship between cell size and external nutrient-imposed growth rate has historically been termed Schaechter’s nutrient growth law^1^. *Escherichia coli* is known to passively correct deviant changes to cell size by adding a constant volume of cellular material (cell unit) between division events, regardless of size at birth^7,8^ which has led to the proposal of what is known as the adder model^9^.

Under steady-state conditions, the cell unit postulated in the adder model is dictated by two mechanisms^10^. The first is an initiation adder that ensures DNA origin firing is triggered only when the cell has grown and accumulated protein factors, such as DnaA, to a threshold required to initiate DNA replication. Synchronous origin firing under conditions of slow growth results in the birth of two daughters, each containing one parental chromosome; however, in environments that promote rapid growth, cell division can occur at a rate faster than the time it takes to duplicate the genome. Bacteria overcome this problem by initiating overlapping replication cycles, so daughter cells are born with two, four or even eight replication origins that ensure genomic integrity is maintained when the cell is faced with rapid division rates^11^. Several studies have reported that the volume of the cell unit proposed in the adder model is proportional to the number of origins of DNA replication present^5,12^, and that the initiation mass coincides with a cell volume per DNA replication origin that is invariable at different growth rates^5,13,14^. Strict regulatory mechanisms ensure timely triggering of replication initiation and co-ordinated progression of elongation and chromosome segregation before cell division occurs^15,16^. The second mechanism of size control is a division adder^10^ that ensures cells only commit to division upon reaching a critical size. It accomplishes this by accumulating factors – such as the division protein FtsZ - to a threshold concentration which allows formation of a Z-ring at midcell along with other essential divisome proteins that eventually trigger septation^17^. Whilst both these methods assist the cell in sensing and controlling its size, the division adder alone allows the cell to correct stochastic deviations in size, ensuring cell size homeostasis^10^.

Cell size, growth and cell cycle progression are known to be influenced by extracellular nutrient abundance. However, the role of intracellular nutrients in these processes is less clear. For example, it is unknown if the major carbon store in bacteria, glycogen, influences cell size. This polyglucan consists of short α-1,4 linked glucan chains joined by α-1,6 branchpoints and three enzymes are known to be involved in its catabolism in the cytosol of *E. coli*. It is initially degraded by glycogen phosphorylase^18^ (GlgP) that removes external glucose moieties, synthesising glucose 1-phosphate (G1P). A glycogen debranching enzyme^19^ (GlgX) cleaves the α-1,6 branchpoints and the resulting soluble malto-oligosaccharides are further mobilised to G1P by GlgP and maltodextrin phosphorylase^20^ (MalP). G1P can then enter pathways involved in central carbon metabolism (CCM), such as glycolysis.

Recent work within our research group generated an *E. coli* Δ*malP/*Δ*glgP/*Δ*glgX* triple mutant strain capable of glycogen synthesis, but severely impaired in its ability to degrade this polymer^21^. In this study we assessed the effect of impairing glycogen breakdown on *E. coli* cell size. We demonstrate that a lesion in *malP* leads to substantial heterogeneity in cell length, revealing a potential link between glycogen turnover and DNA replication.

## RESULTS

### Mutants inhibited in glycogen catabolism grow at the same rate, are equally viable, but show reduced culturability

A synthetic, nutrient-rich medium was employed for our analyses and we examined if growth was affected in isogenic mutant strains lacking *malP, glgP* and/or *glgX*^21,22^ (Table 1). No significant differences in doubling time were observed between the strains (Fig. 1a). We decided to examine the colony forming capacity at three different time points (Fig. 1a) that coincide with early exponential (OD_450_ of ∼ 0.15), onset of stationary (OD_450_ ∼ 4.0) and late stationary phase (OD_450_ ∼ 7.0). During exponential growth, all MalP-deficient strains showed significant reductions in bacterial titre compared to strains with a wild type (WT) *malP* allele (Fig. 1b). During stationary phase, however, none of the mutants differed from the parental strain. Although some differences were detected between the mutant strains during late stationary phase, this was not linked to the presence of a specific mutant allele.

**Table 1.**
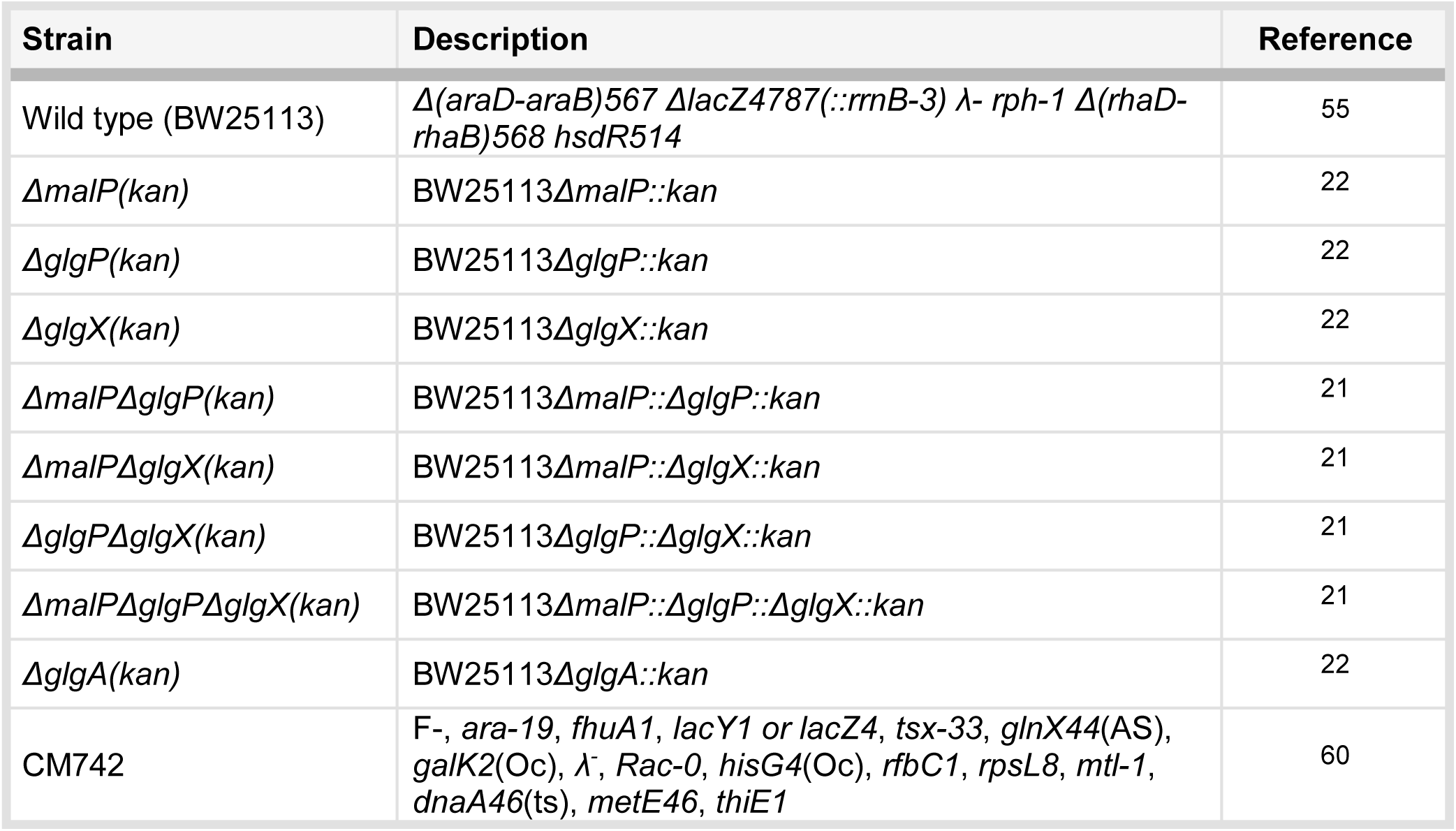
Genotypes of bacterial strains used in the study. Strains were either acquired from the *E. coli* Genetic Stock Center or manufactured in our laboratory.

**Figure 1.**
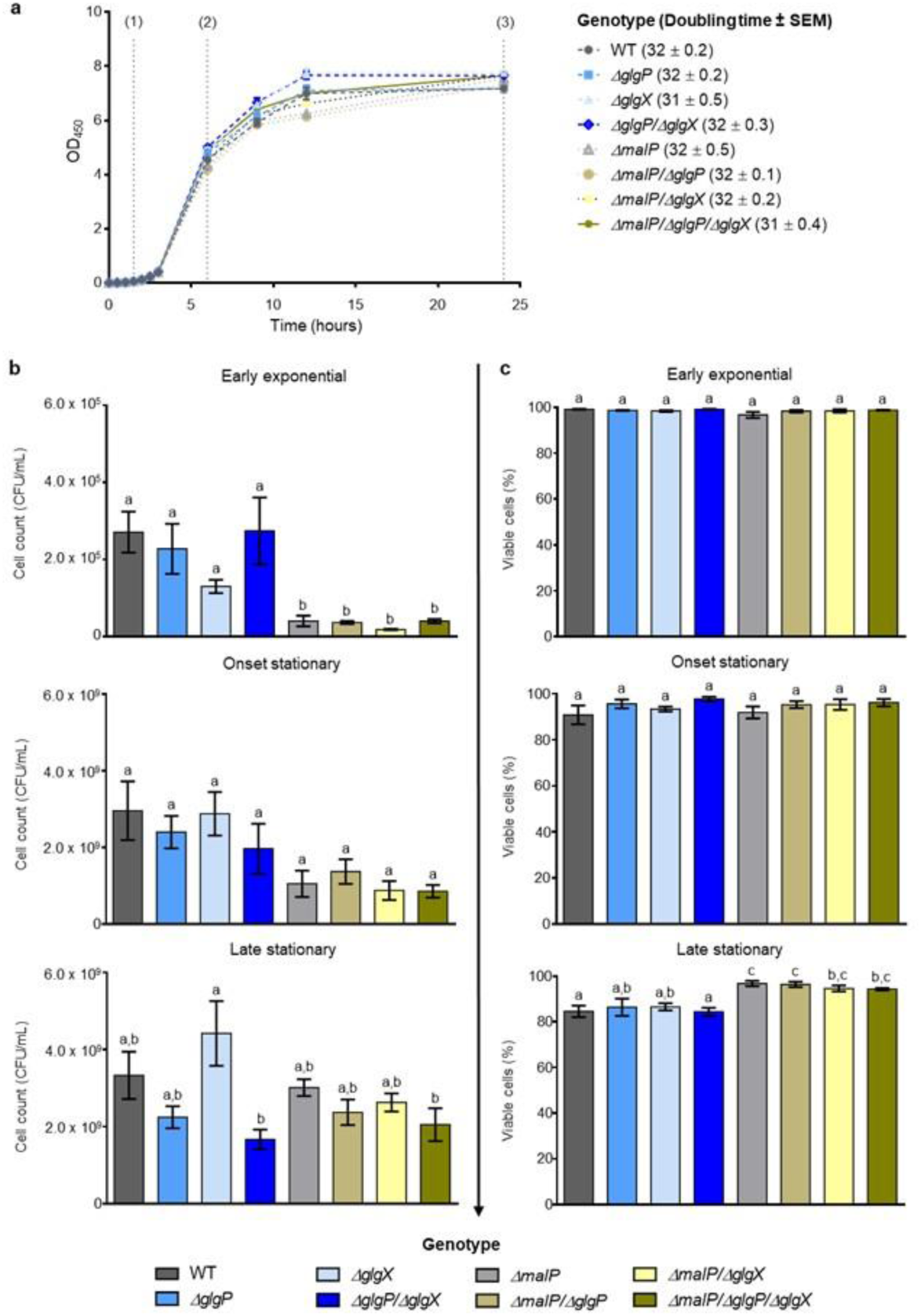
Growth, bacterial culturability and viability. (**a**) Growth curves were generated by determining OD_450_. The mean values from three independent replicates are plotted ± SEM and values within the linear portion of growth were used to calculate doubling times, shown in parentheses in minutes ± SEM. The three time points investigated in this study are indicated by vertical dotted lines and correspond to (1) early exponential growth, (2) onset of stationary phase and (3) late stationary phase. (**b**) Cell counts were ascertained by diluting cells harvested at the indicated times and plating suspensions on LB plates, which were incubated overnight. Colonies were manually scored, and plotted values are the means from three independent replicates ± SEM. (**c**) The percentage of viable cells were determined from confocal images, with plotted values being the means ± SEM from three separate experiments. Each experiment assessed ≥ 550 cells. Different letters in (**b**) and (**c**) denote significant differences (p ≤ 0.05) between group means as determined by a one-way ANOVA with Tukey’s post hoc test.

Cell viability was assessed (Fig. 1c; Supplementary Figs. 1-3) and no differences were observed between any of the strains during exponential growth or at the onset of stationary phase. During late stationary phase, however, mutants with a disrupted *malP* allele produced significantly greater proportion of viable cells than both the parental strain and Δ*glgP/*Δ*glgX*. Furthermore, both Δ*malP* and Δ*malP/*Δ*glgP* formed more viable cells at this stage than either Δ*glgP* or Δ*glgX*, though these single mutants did not differ from either Δ*malP/*Δ*glgX* or the triple mutant.

### All mutant strains produce longer cells than the parental strain during exponential growth

Size parameters of each strain were determined at the same three growth stages using image analysis of bacteria captured by confocal microscopy (Supplementary Figs. 1-3). Width was found to be unaltered between strains (Supplementary Table 1). Individual lengths were sorted into defined intervals to create weighted histograms and difference plots (Supplementary Fig. 4a-c), whilst line charts were generated by grouping lengths into four bins, summarizing the proportion of long cells produced by each strain (Fig. 2a-c). Mutants with a lesion in *malP* demonstrated greater heterogeneity in length during early exponential growth with histograms revealing a marked shift in the prevalence of longer cells, with mean lengths increased from between 2.60-2.82 µm in strains with a WT *malP* allele to between 3.25-3.65 µm in Δ*malP* mutants (Supplementary Fig. 4a).

**Figure 2.**
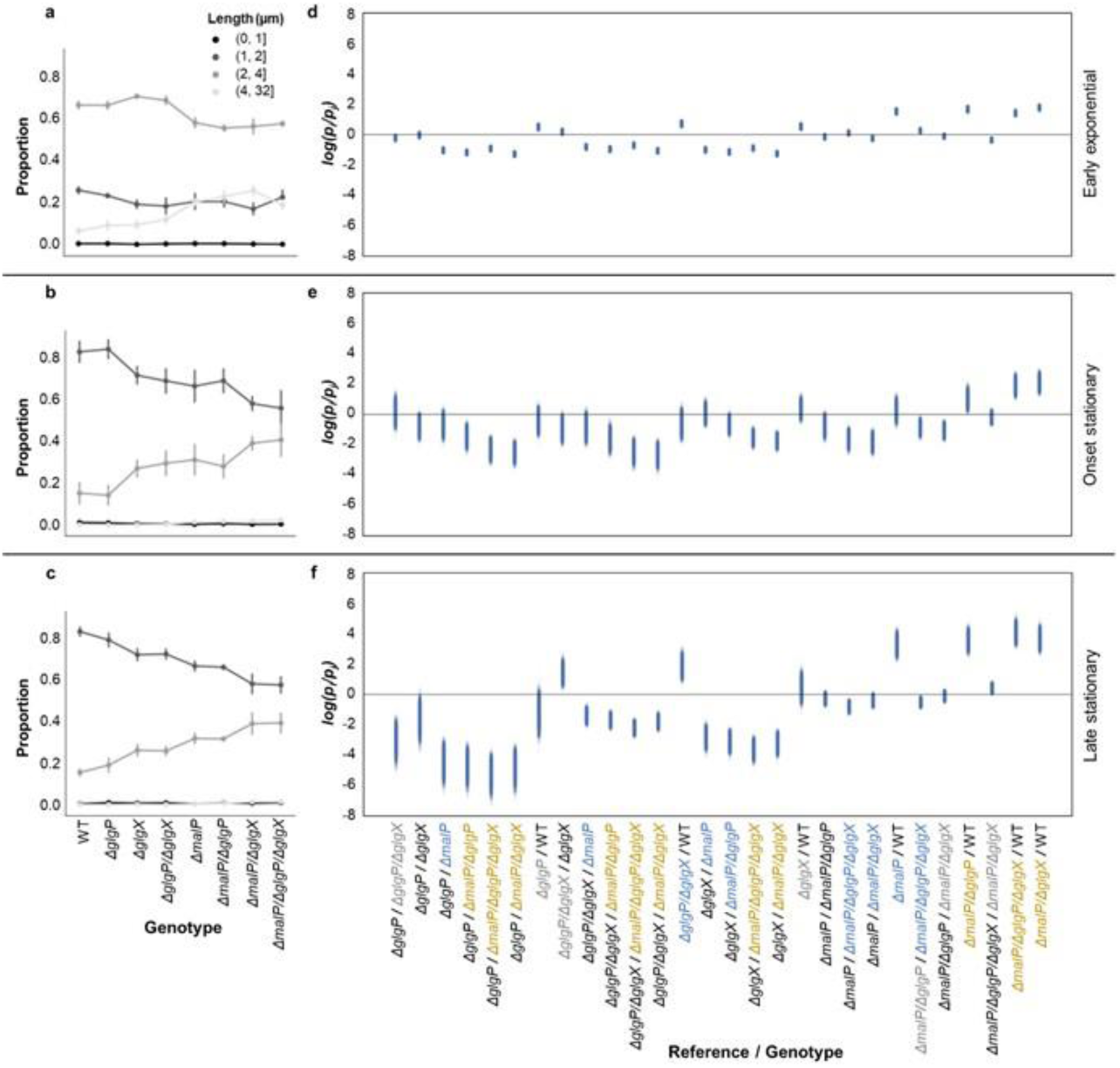
Cell lengths of strains. (**a-c**) Line charts were generated by grouping lengths into four bins according to cut-off points displayed at the upper right corner of the first plot. Data points represent mean ± standard deviation. (**d-f**) The ratios in proportion p of elongated cells (> 4 μm) between reference i and genotype j are denoted p_i_ / p_j_ and scatter plots were created to visualize the spread of p_i_ / p_j_ (see Supplementary Table 2 for the 94% prediction intervals). The further the scatters move away from the baseline, the stronger the evidence that the proportions of elongated cells in reference are different from genotype; if p_i_ = p_j_ then log (p_i_ / p_j_) = 0). Significant increases in elongated cells (p_i_ > p_j_) are coloured grey (at one time point), blue (at two time points) or gold (at all time points). Data were obtained from three independent replicates where ≥ 550 bacteria were measured per experiment.

Variability in cell lengths was greater in Δ*malP* mutant strains, with the standard deviation being over 70% of the mean compared to less than 40% for the other strains during early exponential growth (Supplementary Fig. 4a). Most cells from all strains ranged between 2 to 4 µm in length at this growth phase (Fig. 2a) and the range of cell lengths was also broader here (Supplementary Fig. 4a) than during stationary phase (Supplementary Fig. 4b, c), where all strains contained only a small proportion of cells longer than 4 µm (Fig. 2b, c).

Cells longer than 4 µm were considered elongated. Using Bayesian inference, we examined the probability that each strain accumulates increased numbers of elongated cells by chance and constructed scatter plots to visualize differences between strains (Fig. 2d-f). During early exponential growth all strains with a mutant *malP* allele were significantly enriched in elongated cells, compared to those strains with a WT *malP* allele (Fig. 2d). This was again observed at the onset of stationary phase (Fig. 2e), whilst at the final sampling point the Δ*malP/*Δ*glgX* and Δ*malP/*Δ*glgP/*Δ*glgX* strains formed greater proportions of long cells than all the other strains (Fig. 2f). The Δ*glgP*, Δ*glgX* and Δ*glgP/*Δ*glgX* mutants accumulated more long cells than the WT during exponential growth (Fig. 2d). The Δ*glgP/*Δ*glgX* double mutant differed significantly from the WT, Δ*glgP* and Δ*glgX* strains in its production of elongated cells at the onset of stationary phase, (Fig. 2e), but this was not observed at the final time point (Fig. 2f).

### Z-ring formation and FtsZ amounts are unimpaired in the mutant strains

Membrane-staining revealed invaginating septa at midcell in normal and elongated Δ*malP* mutant cells, along with invaginating septa forming at the ¼ or ¾ cell position in abnormally long bacteria (Supplementary Fig. 1b). To assess whether Z-ring assembly was perturbed in the mutant strains, the percentage of cells with division rings during early exponential growth was determined by FtsZ immunolabelling and confocal microscopy. We found that Z-ring formation was unchanged in the different strains, with ∼70% of cells from all strains showing visible Z-rings during this growth phase (Table 2). Whereas Z-rings were limited to the middle of the cell in bacteria with a WT *malP* allele (Fig. 3a), Δ*malP* mutants formed division rings at midcell in both normal and long cells and at the ¼ and/or ¾ cell position in elongated cells and filaments. Diffuse fluorescent signal was observed in areas between segregated nucleoids in elongated Δ*malP* mutants, suggesting that FtsZ accumulated at putative division sites, but that Z-ring assembly was inhibited. Immunoblots showed that there were no differences in FtsZ levels between any of the strains at any of the time points examined (Fig. 3b, Supplementary Fig. 5).

**Table 2.** Analysis of Z-rings in cells. Confocal images of immunolabeled early exponential phase cells were inspected for the presence of Z-rings. Two independent replicates were performed, assessing ≥ 450 bacteria per experiment.

| Strain | Percentage of cells with a Z-ring (%) |  |
| --- | --- | --- |
|  | Replicate 1 | Replicate 2 |
| Wild type | 70.7 | 68.4 |
| $\Delta glgP$ | 69.8 | 65.9 |
| $\Delta glgX$ | 67.5 | 66.3 |
| $\Delta glgP/\Delta glgX$ | 67.8 | 64.2 |
| $\Delta malP$ | 72.7 | 67.1 |
| $\Delta malP/\Delta glgP$ | 73.2 | 67.4 |
| $\Delta malP/\Delta glgX$ | 67.7 | 67.5 |
| $\Delta malP/\Delta glgP/\Delta glgX$ | 67.8 | 67.4 |

**Figure 3.**
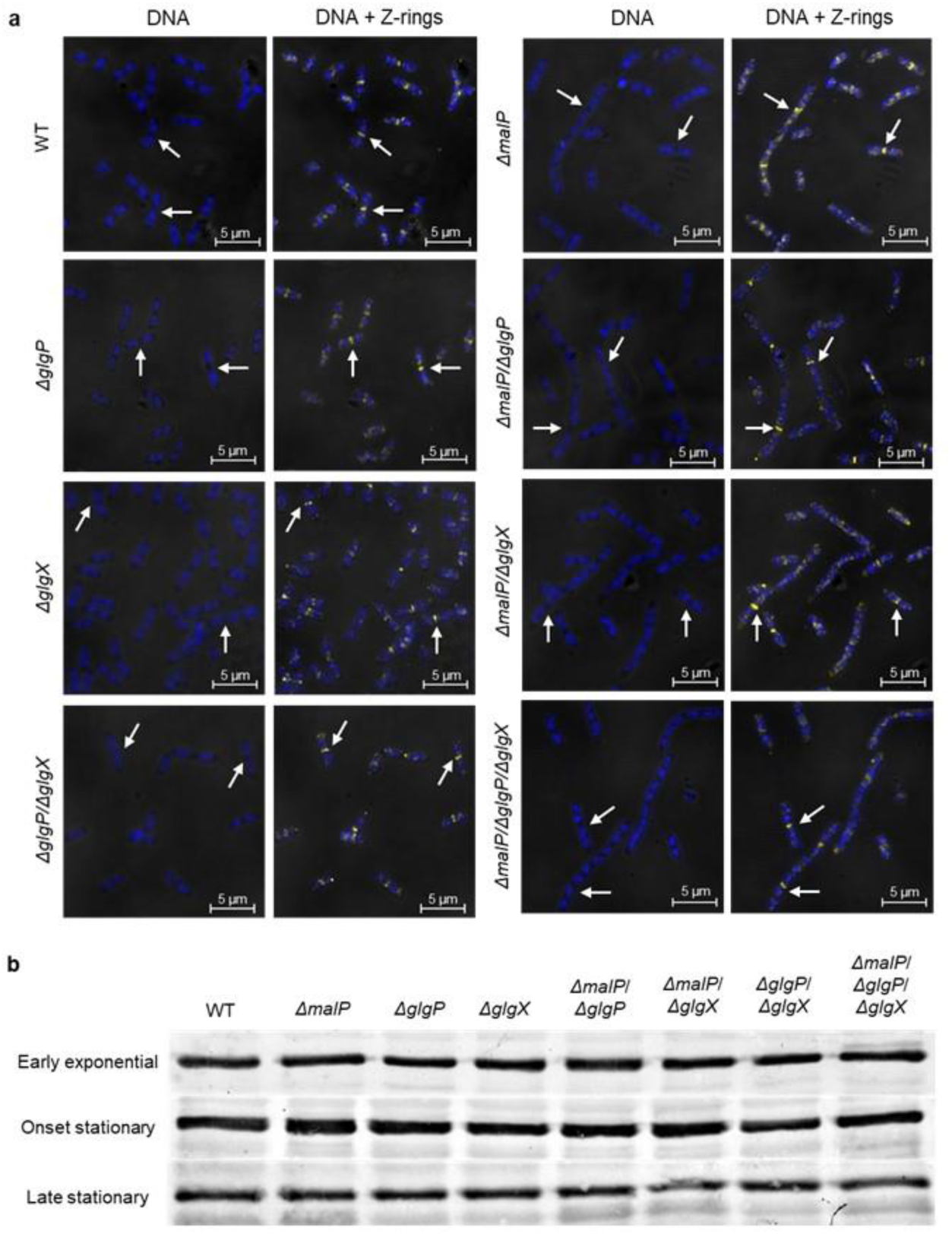
Z-ring formation and FtsZ amounts. (**a**) Bacteria harvested at OD_450_ of ∼ 0.15 were fixed and immunolabelled using an α-FtsZ antibody. Phase contrast and confocal microscopy images depict the positions of Z-rings in cells during early exponential growth. These structures are shown in yellow, whilst DAPI-stained nucleoids are displayed in blue. White arrows indicate the positions of Z-rings. (**b**) Immunoblot analysis of relative FtsZ levels. Ten micrograms of total crude protein were separated by 10% (w/v) SDS-PAGE and visualized following labelling with an anti-FtsZ antibody.

### Glycogen and protein aggregates are observed in Δ*glgX* mutants

Glycogen amounts increased in all the strains upon entry into stationary phase, with cells in late stationary phase containing the most (Fig. 4a). The triple mutant accumulated significantly more glycogen than the WT both at the onset (p = 0.026) and in late (p = 0.038) stationary phase, although these were the only significant differences detected at any time point.

**Figure 4.**
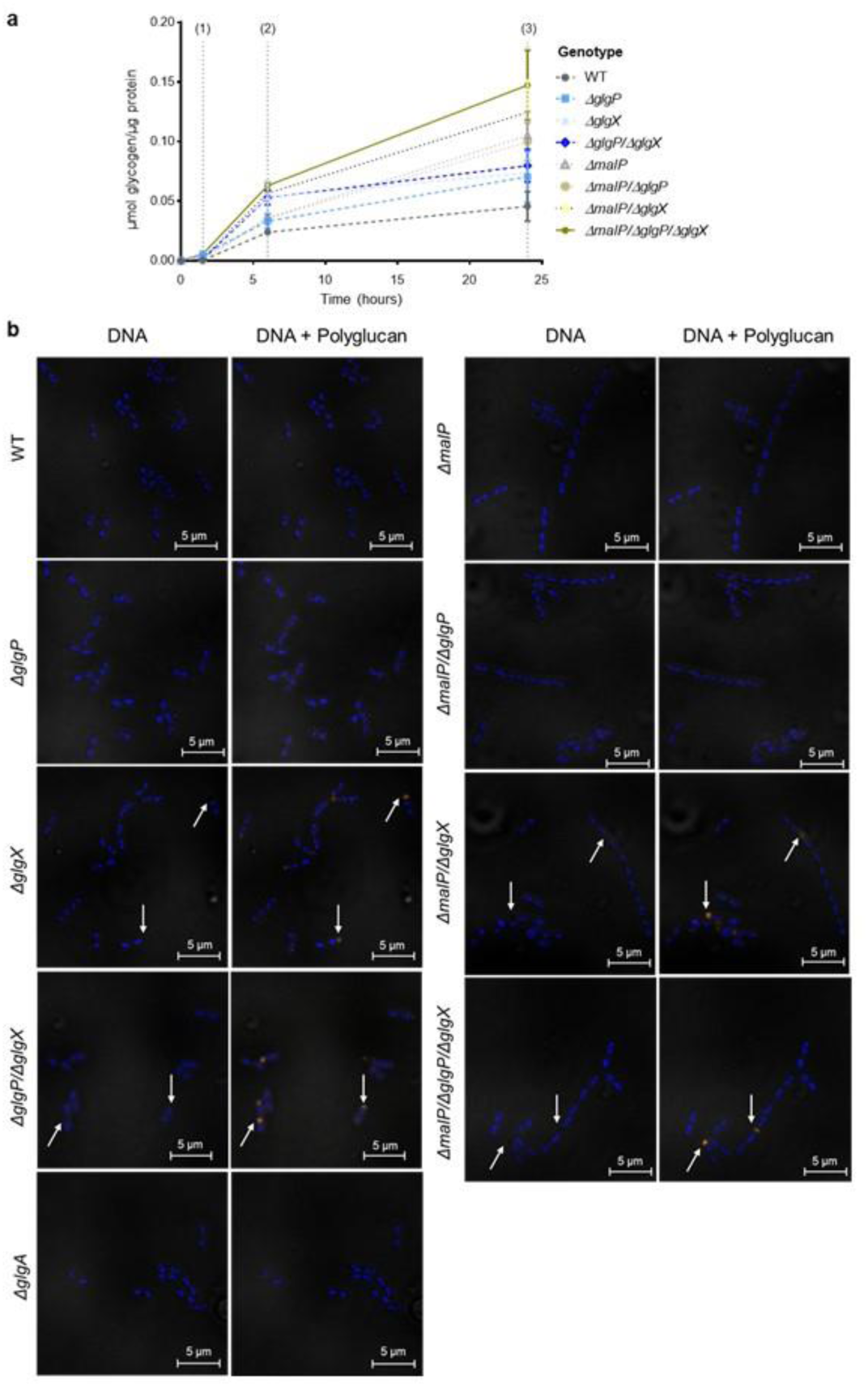
Glycogen amounts staining of glycogen and DNA in exponentially growing bacteria. (**a**) Glycogen concentrations were determined at the three indicated time points via an enzyme-linked assay. Values plotted are means from three independent repeats ± SEM. (**b**) Cells were cultured to early exponential phase and then harvested, fixed and stained for polyglucan and DNA before being imaged by confocal and phase contrast microscopy. DAPI-stained DNA is shown in blue and PAS-stained glycogen in orange. White arrows indicate the positions of glycogen bodies.

We visualized glycogen aggregates in the strains as well as in a glycogen synthase (Δ*glgA*) mutant^22^. As expected^23^, no staining was observed in the Δ*glgA* strain (Fig. 4b; Supplementary Figs. 6-7) demonstrating the specificity of the stain. We observed substantial glycogen bodies in all strains with a defective *glgX* allele during early exponential growth (Fig. 4b). Whilst polar glycogen aggregates were observed in these strains, glycogen bodies were also noted in nucleoid-free regions and, infrequently, at midcell within elongated bacteria where both *malP* and *glgX* are mutated (Δ*malP/*Δ*glgX* and Δ*malP/*Δ*glgP/*Δ*glgX*). Polar polyglucan bodies appeared in all strains upon entry into stationary phase, with some diffuse signal detected along the cell peripheries (Supplementary Fig. 6). Glycogen bodies were particularly pronounced in strains with a defective *glgX* allele and were again detected at inter-nucleoid zones within elongated Δ*malP/*Δ*glgX* and Δ*malP/*Δ*glgP/*Δ*glgX* cells. This phenotype was less apparent in all strains cultured to late stationary phase (Supplementary Fig. 7) as staining was restricted to the cell poles and periphery.

Protein aggregates, stained by the amine-reactive viability dye, were observed only weakly in GlgX-deficient strains during early exponential growth, and their spatial arrangement at midcell and polar regions coincides with that of glycogen bodies (Supplementary Fig. 1). At later growth stages, protein granules of various sizes were observed mainly at the poles of all strains (Supplementary Figs. 2-3).

### Strains lacking MalP produce multinucleate filamentous cells, but there is no alteration in numbers of anucleate cells

During early exponential growth, DNA staining demonstrated no obvious phenotypic differences between Δ*glgP*, Δ*glgX* and Δ*glgP/*Δ*glgX* mutants and the WT (Figs. 3a, 4b; Supplementary Fig. 1a). Genetic material within elongated cells from Δ*malP* mutants appeared as unsegregated masses or as regularly spaced nucleoids along the length of the cell (Figs. 3a,4b; Supplementary Fig. 1b). During stationary phase MalP-deficient mutants again demonstrated aberrant nucleoid partitioning, characterized by clumps of DNA (Supplementary Figs. 2b, 3b, 6, 7), whilst this was not observed in the other strains (Supplementary Figs. 2a, 3a, 6, 7).

We determined the percentage of achromosomal cells formed by the strains over time and observed no significant differences in anucleate cell production between any of the strains during either exponential growth, or at the onset of stationary phase (Fig. 5a). In late stationary phase, the Δ*glgX* mutant produced significantly fewer anucleate cells than the WT (p = 0.035) but no other differences were detected between groups.

**Figure 5.**
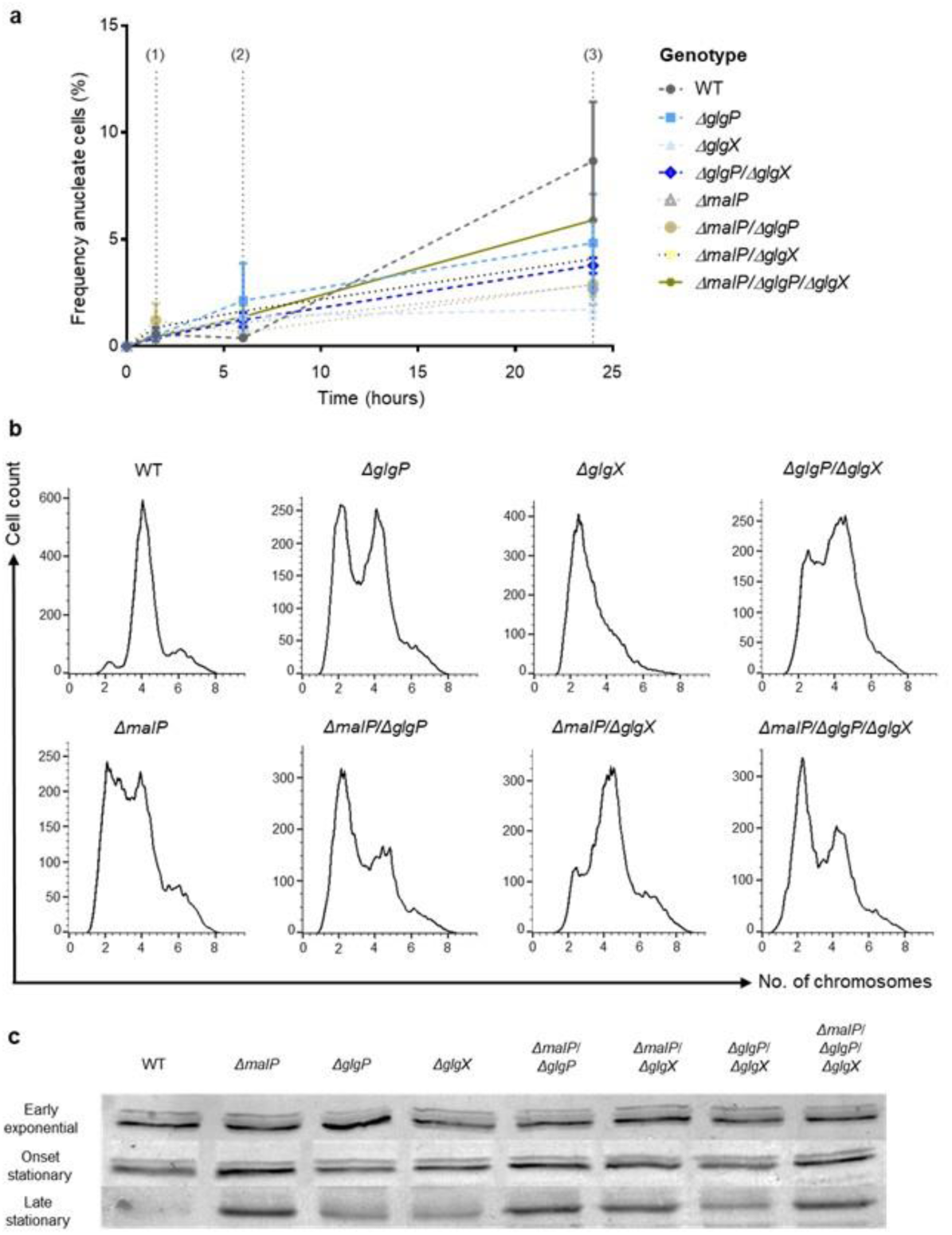
Achromosomal cell production, replication patterns and DnaA levels. (**a**) Confocal pictures of DAPI-stained bacteria were assessed at the three indicated time points to determine the percentage of achromosomal cells. Data represents the mean counts ± SEM from three independent experiments evaluating ≥ 550 cells per repeat. (**b**) Drug-treated, early exponential phase (OD_450_ ∼ 0.15) bacteria were stained with Hoechst, analyzed by flow cytometry and histograms depict the DNA content of such cells. The y-axis represents cell count per channel and the x-axis chromosome number. Diagrams depict the outcome of a single experiment where ∼ 20 000 events were captured. (**c**) Immunoblot examination of relative DnaA levels. Each lane was loaded with 10 µg total protein and separated by 10% (w/v) SDS-PAGE, before being blotted onto nitrocellulose membranes and visualized following probing with an α-DnaA antibody.

### Mutant strains display diverse replication patterns

DNA concentrations relative to the WT (Table 3) were assessed during early exponential growth. The DNA and protein contents of individual cells were measured by flow cytometry. Total protein of an *E. coli* cell may be used as a measure of its mass^24^. The DNA content of the Δ*malP* mutant was more than 40% higher than the WT, but its DNA concentration (DNA/mass) was unchanged due to a concomitant increase of nearly 40% in its mass (Table 3). The other MalP-deficient mutants and the Δ*glgP/*Δ*glgX* strain had cell mass values that were 9-20% higher than the WT. The DNA concentration of strains with a mutant Δ*glgP* and Δ*glgX* allele (Δ*glgP/*Δ*glgX* and Δ*malP/*Δ*glgP/*Δ*glgX*) were reduced by more than 10%, relative to the WT (Table 3).

**Table 3.** DNA amounts and cell mass. Mean DNA content, cell mass and DNA/mass of untreated mutant *E. coli* strains relative to the wild type during early exponential growth. Data represent means ± standard deviation calculated from four independent replicates surveying ∼ 20 000 cells per run.

| Strain | Relative DNA content | Relative mass | DNA/mass |
| --- | --- | --- | --- |
| Wild type | 1.00 ± 0.4 | 1.00 ± 0.4 | 1.00 |
| *ΔglgP* | 1.04 ± 0.4 | 1.01 ± 0.4 | 1.04 |
| *ΔglgX* | 1.02 ± 0.4 | 1.05 ± 0.4 | 0.98 |
| *ΔglgP/ΔglgX* | 1.00 ± 0.4 | 1.18 ± 0.4 | 0.85 |
| *ΔmalP* | 1.48 ± 0.4 | 1.37 ± 0.4 | 1.08 |
| *ΔmalP/ΔglgP* | 1.08 ± 0.4 | 1.09 ± 0.4 | 0.99 |
| *ΔmalP/ΔglgX* | 1.06 ± 0.4 | 1.16 ± 0.4 | 0.91 |
| *ΔmalP/ΔglgP/ΔglgX* | 1.06 ± 0.5 | 1.21 ± 0.4 | 0.86 |

Early exponential phase cells were treated with rifampicin and cephalexin which prevents the initiation of new rounds of DNA replication^25^ and blocks cell division^26^, respectively. Cells incubated under these run-out conditions end up with an integer number of chromosomes, reflecting the number of origins present in a cell at the time of antibiotic addition. DNA replication patterns of the different strains were subsequently analysed by flow cytometry (Fig. 5b). In the WT strain, DNA resolved as three distinct peaks correlating to 2, 4 or 6 chromosome equivalents. The largest proportion of initiated cells contained 4 origins, indicating synchronous origin firing, though the presence of 6 origin cells suggests a degree of replication asynchrony under our experimental conditions. The peak of cells with 2 chromosomes represents bacteria that had not initiated replication when antibiotics were added, and a delay in initiation would increase the proportion of such cells in a population^27^. Such a delay is observed in most of the mutants whilst problems with completing ongoing replication are apparent from poorly resolved peaks^28–30^.

Mutant Δ*glgP* cells showed two distinct peaks corresponding to cells with 2 or 4 chromosomes, though these did not separate completely, suggesting the existence of a subpopulation of cells containing non-integer chromosome numbers between 2 and 4 chromosomes. There appeared to be a considerable initiation delay in this strain, indicated by the presence of a large proportion of cells with 2 origins. The presence of bacteria containing between 2 and 4 origins probably indicates that replication forks could not reach the termini, despite the length of this assay (3 hours). Initiation is also clearly delayed in the Δ*glgX* mutant, though the broadness of the peak suggests that existing replication cycles could not be completed in this strain and thus denotes cells with at least 2 and 4 origins.

The replication pattern in the Δ*glgP/*Δ*glgX* strain appears to be intermediate between the Δ*glgP* and Δ*glgX* single mutants where initiation is delayed, and elongation perturbed. Cells with at least 2 and 4 origins are present, though peaks are poorly separated and not as distinct as in the Δ*glgP* single mutant, indicating that disruption of *glgX* possibly has an additional negative effect on replication fork progression in this strain. Furthermore, the Δ*malP* single mutant showed a similar broad and undefined peak of cells with 2 to 4 origins, though cells containing up to 6 chromosomes are also present.

The severe replication problem observed in Δ*malP* single mutant cells is partially alleviated when *glgX* is also mutated. An initiation delay is still apparent from the substantial proportion of 2 chromosome cells, though the appearance of a distinct subpopulation of 4 origin cells in the Δ*malP/*Δ*glgX* strain implies that origin firing more closely resembles that of the WT than either of the single mutants (Δ*malP* or Δ*glgX*). Problems with stalled forks were less severe in this double mutant than the Δ*malP* single mutant, though cells with partially replicated chromosomes are still apparent, containing between 2 and 4 or 4 and 6 genome equivalents. Disruptions to replication fork progression seen in the Δ*malP* single mutant also appear to be ameliorated by delaying initiation upon elimination of *glgP* in either a Δ*malP* or Δ*malP/*Δ*glgX* mutant background. Both Δ*malP/*Δ*glgP* and the triple mutant produce identical replication patterns with large proportions of uninitiated, 2 origin cells. Bacteria with 4 origins are also present along with those where replication could not be completed, and such cells thus contain between 2 and 4 chromosomes.

Immunoblots revealed no marked differences in DnaA levels between any of the strains during exponential growth (Fig. 5c; Supplementary Fig. 8), though levels of this initiator protein were elevated in Δ*malP* mutants upon entry into stationary phase with these differences becoming greater by late stationary phase.

## DISCUSSION

One of the major carbon stores in bacteria is glycogen and its breakdown is linked to CCM. We decided to examine if glycogen degradation influences bacterial cell size using a set of mutants impaired in their ability to degrade this polyglucan. This was due to the observation that, although the cells from all strains grew at the same rate (Fig. 1a), the number of culturable cells in strains containing a mutant *malP* allele were reduced during exponential phase (Fig. 1b). Fewer culturable cells present at the same optical density could indicate that bacteria were forming non-viable or anucleate cells, or were growing larger without dividing, and our data support the latter explanation.

During early exponential growth, all strains accumulated cells which were longer than those found during stationary phase (Fig. 2a-c; Supplementary Fig. 4), and strains with a mutant *malP* allele were significantly enriched in elongated cells (Fig. 2d) that included a small proportion of filaments longer than 4 µm (Fig. 2a). Cell mass values of all MalP-deficient strains (Table 3), relative to the WT, were marginally elevated during exponential growth, consistent with the finding that these strains contained significant proportions of elongated cells at this growth phase (Fig. 2d). Invaginating septa were observed forming at midcell or one cellular unit away from a cell pole in elongated, exponentially growing MalP-deficient bacteria (Supplementary Fig. 1b). No detectable differences in viability (Fig. 1c) or anucleate cell formation (Fig. 5a) were found between strains at this growth phase, which indicates that the reason Δ*malP* mutants show reduced culturability during early exponential growth (Fig. 1b) is partially caused by the formation of elongated and filamentous cells.

It is well-known that delaying cell cycle progression alters cell length. For example, arresting initiation of DNA replication^10,31^ or delaying the onset of septation^12^ have been shown to result in significant increases in cell length, whilst width is either unchanged or reduced^32^. On the other hand, disruptions to fatty acid metabolism alter cell length, but not width^33,34^. In our experiments cell width remained invariable (Supplementary Table 1) indicating that the observed alteration in cell length is most likely the result of some downstream effect on the cell cycle, rather than on fatty acid metabolism.

Mutant Δ*malP* strains showed significant heterogeneity and enhanced lengths during early exponential growth (Fig. 2a, d; Supplementary Figs. 1b, 4a) whilst mass doubling times were unchanged between strains (Fig. 1a). Genetic alterations that lead to a contravention of Schaechter’s nutrient growth law exist, including moderate delays in division that have no observable effect on mass doubling times, but enhance cellular lengths^12,35^. It appears that elongated Δ*malP* mutants experience a partial defect in cell division, though this is unlikely due to differences in FtsZ levels (Fig. 3b) or Z-ring formation (Table 2) as these parameters were identical between strains. Furthermore, MalP-deficient cells clearly showed nucleoid partitioning defects during exponential growth (Figs. 3a, 4b; Supplementary Fig. 1b), which is expected to delay cell constriction. This suggests that maternal cells experience difficulties coordinating DNA replication with nutrient availability, cell size and mass doubling. Stochastic variabilities in length likely results from bacteria that are unable to control synchronous progression of cell cycle events^32,34,36^ – a situation which has been reported in other mutants affected in CCM^32^. Hence, we investigated the cell cycle.

In filamentous cells, size homeostasis is ensured by employing adder-like mechanisms alongside the Min system to sense absolute size and direct division rings to form one cell unit away from the pole, and two from another ring^36^. This allows progeny of even the longest cells to converge at a population average after multiple generations, as was observed in the MalP-deficient strains by late stationary phase (Supplementary Fig. 4). Given the lack of alteration in division plate numbers (Table 2), it is unsurprising that FtsZ amounts were similar in all strains at all growth stages (Fig. 3b). Taken together this demonstrates that the impairment of division is not caused by decreased capacity to form division rings but is instead the result of some mechanism(s) delaying septation.

Elongated, early exponential phase MalP-deficient cells contain multiple nucleoids which include masses of genetic material that are poorly separated (Figs. 3a, 4b; Supplementary Fig. 1b). Unsegregated clumps of DNA are also evident in these strains during stationary phase (Supplementary Figs. 2b, 3b, 6, 7). Run-out histograms indicated that mutations affecting *glgP* seem to stall replication initiation irrespective of genotype (Fig. 5b). Cells from the Δ*malP* single mutant display aborted replication forks and do not resolve into distinct populations containing integral number of chromosomes (Fig. 5b). Double and triple mutants with lesions in *glgP* and *malP* produce similar run-out histograms (Fig. 5b). This indicates that mutating *glgP* delays replication initiation and, therefore, likely reduces replication rates in both the Δ*malP* and Δ*malP/*Δ*glgX* mutant backgrounds. Deleting *glgX* alone delayed replication initiation in the single mutant, while this effect appears masked when it is mutated alongside *glgP* (i.e. Δ*glgP/*Δ*glgX* and Δ*malP/*Δ*glgP/*Δ*glgX* strains). The replication patterns of a double Δ*malP/*Δ*glgX* mutant closely resembles that of the WT. It is possible that disrupting *glgX* in the Δ*malP* single mutant delays replication and affords the cell more time to overcome difficulties with stalled forks replication (Fig. 5b) or nucleoid segregation (Figs. 3a, 4b; Supplementary Fig. 1b).

Increasing DnaA concentrations in *E. coli* has been shown to cause over-initiation and filamentation with replication fork stalling observed under run-out conditions and flow cytometry examination^37^, however, we observed no differences in DnaA amounts between strains during early exponential growth when replication patterns and DNA amounts were assessed (Fig. 5c). This indicates that alterations in exponential phase DnaA levels do not provide an explanation for the observed fork stalling in our experiments. The cause and implication of elevated stationary phase DnaA levels on DNA replication in Δ*malP* mutants remains unclear.

Length increases were not exclusive to cells from strains with a mutated *malP* allele as Δ*glgP*, Δ*glgX* and Δ*glgP/*Δ*glgX* mutants formed greater proportions of longer cells more frequently than the WT during exponential growth (Fig. 2d). While disrupting the normal progression of DNA replication (Fig. 5b) sufficiently explains altered cell lengths in these mutant strains (Fig. 2d), it is uncertain whether this is the entire picture. Discrepancies in the proportion of elongated cells produced by mutant strains with a lesion in Δ*glgX* (especially in combination with a mutant *malP* allele) became more pronounced during stationary phase (Fig. 2b, c; Supplementary Fig. 4b, c). This is typified by the observation that Δ*malP/*Δ*glgX* and the Δ*malP/*Δ*glgP/*Δ*glgX* strains accumulate enhanced proportions of long cells compared to the Δ*malP* and Δ*malP/*Δ*glgP* strains by late stationary phase (Fig. 2c, f).

Since glycogen over-accumulation has been associated with cell size dysregulation upon entry into stationary phase in *E. coli csrA* mutants^38^, we investigated whether our mutant strains accumulated excessive amounts of this biopolymer. Disruptions to *glgP* and/or *glgX* have been reported to increase glycogen concentration as turnover rates are diminished^18,19,21,39^. Previously, we observed significant increases in stationary phase glycogen amounts in all GlgX-deficient strains cultured in LB broth with glucose^21^. However, the only difference we could detect using the current nutritional regime was between the WT and triple mutant during stationary phase (Fig. 4a). The reason for this discrepancy is most likely due to differences in media composition that are known to affect bacterial glycogen accumulation^40^.

Glycogen deposits were observed at polar sites in all strains at the onset of stationary phase. This was most clear in Δ*glgX* mutant strains and deposits also formed at inter-nucleoid zones when *malP* was concurrently disrupted (Supplementary Fig. 6). Most cells harvested during late stationary phase demonstrated a combination of polar and peripheral PAS staining (Supplementary Fig. 7). Despite the low amounts of glycogen detected during exponential growth (Fig. 4a) we were able to visualize glycogen bodies in all strains containing a lesion in *glgX* (Fig. 4b), again appearing at inter-nucleoid zones and potential division sites (Fig. 3a) and not only at the cell periphery as reported previously for *E. coli*^41–43^. This indicates that small amounts of glycogen remain within these cells upon exiting lag phase^44,45^, possibly due to structural changes that are known to increase short chains within this polymer^19,21,39^ and which would be expected to delay turnover^39^.

Protein inclusion bodies were observed from viability micrographs in GlgX-deficient strains during exponential growth (Supplementary Fig. 1), and their occurrence at polar or inter-nucleoid sites mimics the spatial distribution of glycogen bodies (Fig. 4b). In *E. coli*, sites of protein aggregation are spatially dictated by nucleoid occlusion^46^, where the bacterial nucleoid sets a zone of high macromolecular crowding around itself. Damaged proteins are passively sequestered to these sites^46,47^, effectively restricting protein inclusion bodies to DNA-free regions within the cytosol. Excess glycogen, like polypeptides, are likely targeted passively to sites of low macromolecular crowding, determined by nucleoid occlusion^46^, following synthesis at the cell periphery. Within the context of endogenous metabolism, it is probable that such sequestration mechanisms protect macromolecules, like proteins, from turnover, as the cell will preferentially degrade glycogen for carbon and energy^48,49^.

Coinciding glycogen and protein aggregates could explain why DNA concentrations are reduced and cell mass values (protein content) elevated in strains where *glgP* and *glgX* are concurrently disrupted (Table 3) as these two strains (Δ*glgP/*Δ*glgX* and Δ*malP/*Δ*glgP/*Δ*glgX*) showed the most conspicuous glycogen (Fig. 4b) and protein inclusion bodies (Supplementary Fig. 1) during exponential growth. At the onset of stationary phase, the Δ*glgP/*Δ*glgX* double mutant, which displayed prominent glycogen deposits (Supplementary Fig. 6) was significantly enriched in long cells compared to the WT, Δ*glgP* and Δ*glgX* strains (Fig. 2e). Together with the observation of distinct glycogen bodies in the Δ*malP/*Δ*glgX* and Δ*malP/*Δ*glgP/*Δ*glgX* strains at the onset of stationary phase (Supplementary Fig. 6), and the finding that these two strains produced significantly more long cells than the Δ*malP* and Δ*malP/*Δ*glgP* strains at the final sampling point (Fig. 2c, f), it is plausible that glycogen deposits may additionally increase cell lengths.

Our results provide novel insight into how cell lengths are altered in mutants affecting glycogen degradation. DNA replication is clearly affected in the mutant strains. Whether this results from disturbing carbon flux and altering energy levels, or the levels of specific metabolites that synchronize cell cycle events is unclear. Disruptions to DNA replication may be the result of currently unidentified protein-protein interactions between enzymes involved in glycogen catabolism or CCM and those involved in controlling certain aspects of the cell cycle. A high-throughput study of protein-protein interactions in *E. coli* revealed that MalP interacts with diverse proteins known to regulate the SOS response, DNA damage repair and metabolism ^50^. Links between CCM and DNA replication have been demonstrated previously^51–53^ though how MalP helps regulate this fundamental process remains unclear. Phosphorylases produce substantial amounts of G1P during glycogen degradation, which may be fed into CCM after being converted to G6P. Disruptions to CCM and UDPG levels are known to affect cell size^32,54^, so it is tempting to speculate that these may be affected pleiotropically in the mutants in this study and our future work will examine these hypotheses.

## MATERIALS AND METHODS

### Strains affected in glycogen degradation and growth analysis

All strains (Table 1) affected in glycogen metabolism are derivatives of the BW25113^55^ and form part of the isogenic Keio collection^22^. They were obtained either from the *E. coli* Genetic Stock Centre (CGSC, Yale University) or were manufactured as described previously^21^.

Cells were cultured at 37°C in AB medium^56^ supplemented with 10 µg/mL thiamine, 25 µg/mL uridine^57^, 0.4% (w/v) glucose and 0.5% (w/v) casamino acids (ABTGCU medium). The slowly growing BW25113 cells, used as a standard for flow cytometry, were cultured in the same minimal medium, except that uridine and casamino acids were omitted (ABTG medium). To achieve balanced growth, cells were grown overnight from a single colony and then diluted 1:1000 into fresh medium. Bacteria were cultured until an OD_450_ ∼ 0.2 - 0.4 was reached, at which point cells were back-diluted to an OD_450_ ∼ 0.005 into fresh medium. Growth was followed spectrophotometrically at 450 nm at 30-minute intervals for the first three hours of growth and doubling times calculated from OD values using an online tool (http://www.doubling-time.com/compute_more.php). Colony forming units were assessed by serially diluting an aliquot of cells in GTE (50 mM Glucose, 25 mM Tris-HCl (pH 8.0), 10 mM EDTA) and plating 10 µL on LB agar plates. Plates were incubated for 12 hours at 37°C, at which point colonies were counted manually.

### Glycogen and protein quantification

Duplicate samples of either 45 mL (exponential phase cells) or 2 mL (stationary phase cells) were harvested by centrifugation (20 000 x g, 10 minutes, 4°C) and cell pellets frozen at −80°C until glycogen levels were determined using a previously published method^21^.

For determination of total protein, cell pellets were re-suspended in an equal volume of protein extraction buffer (PEB, 17 mM Tris-HCl pH 8.5, 5 mM EDTA, 1 mM phenylmethylsulfonyl fluoride (PMSF) and 2 mM β-mercaptoethanol) and lysed through sonication (10 bursts lasting 15 seconds each, at 5 W output, with 1-minute cooling between bursts). Cell suspensions were clarified by centrifugation (20 000 x g, 10 minutes, 4°C) and total soluble protein quantified^58^ using a kit (Bio-Rad Protein Assay Kit) and BSA as a standard.

### Immunoblotting

Cells were harvested, washed once in TE buffer (20 mM Tris-HCl pH 7.5, 1 mM EDTA) and re-suspended in PEB, before being lysed by sonication and using soluble protein concentrations determined as described above. Samples containing 10 µg total protein were separated by 10% (w/v) SDS-PAGE and transferred onto a PVDF membrane by semi-dry blotting using standard procedure^59^. Membranes were probed either with α-FtsZ (1:2000 dilution) serum (Agrisera) or α-DnaA (1:1500 dilution) serum followed by goat anti-rabbit secondary antibody, conjugated to alkaline phosphatase. Immunoblots were developed by incubating in BCIP/NBT solution (Sigma) for 10 minutes before imaging.

A mutant *dnaA46* strain (CM742^60^; Table 1), was used as a negative control in the blots assessing DnaA protein amounts. This strain was cultured at 30°C in LB broth until early exponential phase (OD_600 ∼_ 0.1 – 0.2) and then shifted to 42°C (non-permissive temperature). Growth continued for 1.5 hours at which point cells were harvested and soluble proteins extracted and quantified as described above.

### Flow cytometry

Cells in early exponential phase (OD_450 ∼_ 0.1 – 0.2) were either directly harvested by centrifugation (3 300 x g, 5 minutes, 4°C) or treated with 150 µg/mL of rifampicin (USBiological) and 45 µg/mL of cephalexin (Sigma) for at least 3 hours for replication run-out experiments. Drug-treated cells were subsequently collected by centrifugation and both treated and untreated cells re-suspended in ice-cold TE buffer before being fixed by diluting tenfold in 77% (v/v) ice-cold ethanol^61^.

Samples were kept at 4°C for at least 12 hours before being pelleted by centrifugation and air dried to remove any residual ethanol^62^. Untreated samples used for ratiometric analyses were washed once in 0.1 M phosphate buffer (pH 9.0) and re-suspended in the same buffer containing fluorescein isothiocyanate (FITC) at a final concentration of 5 µg/mL to stain total cellular protein^63^. After overnight staining at 4°C, cells were washed twice in TBS (20 mM Tris-HCl pH 7.5, 130 mM NaCl) and genetic content stained with 1.5 µg/mL of Hoechst 33258 (Sigma) in the same buffer^64^. Run-out samples were similarly washed in TBS and DNA stained with Hoechst in the same buffer. Hoechst staining proceeded for at least 30 minutes on ice, prior to flow analysis.

Flow cytometry was performed using a BD FACS Aria™ IIu, equipped with a violet (407 nm) and blue (488 nm) laser, and data recorded using BD FACSDIVA™ 8.0.1 software. Hoechst 33258 fluorescence was captured using a 450/40 bandpass filter, whilst FITC fluorescence was collected through a 530/30 bandpass filter. Average cell mass was taken as median FITC fluorescence, whilst average DNA content per cell was determined as median Hoechst fluorescence. The relative DNA per mass (DNA concentration) (ratio) was calculated by dividing average DNA content by average cell mass^65^. As a control for chromosome number, an ethanol-fixed sample of untreated, slowly growing BW25113 cells containing mainly 2 chromosomes was run as an external standard. Data obtained from flow experiments were processed by FlowJo v10.6.1 (©Tree Star, Inc) software.

### Membrane and vital staining techniques

An aliquot of cells corresponding to ∼ 10^5^ CFU/mL were collected at the indicated growth stages by centrifugation (3 300 x g, 5 min, 22°C) and re-suspended in 1 mL 1X PBS (137 mM NaCl, 2.68 mM KCl, 10.1 mM Na_2_HPO_4_, 1.76 mM KH_2_PO_4_). Cell envelopes were stained by adding 5 µL of 1 mg/mL FM™ 4-64FX (Thermo Fisher Scientific) and non-viable cells stained with 1 µL of the LIVE/DEAD™ cell dye (Thermo Fisher Scientific). Samples were incubated at room temperature for 5 minutes and then fixed using an established method with slight modifications^66^, through the addition of 200 µL ice-cold fixative (16% (v/v) formaldehyde, 0.06% (v/v) glutaraldehyde in 1X PBS). Following incubation at room temperature for 15 minutes with slight agitation (100 RPM), cells were kept on ice for another 30 minutes before being collected by centrifugation (20 000 x g, 1 minute, 4°C) and washed three times with 1X PBS containing 1 mM EDTA (PBS-E). Bacteria were re-suspended in GTE and stored at 4°C. Mounting and imaging occurred within 48 hours of sample preparation.

### Periodic acid-Schiff (PAS) staining

An aliquot of cells corresponding to ∼ 10^5^ CFU/mL were collected by centrifugation (3 300 x g, 5 min, 22°C) and re-suspended in 1 mL PBS. Cells were fixed essentially as described above, except that glutaraldehyde was omitted from the fixative^67^. Following the third wash step with PBS-E, cells were instead collected and resuspended in 100 µL ice-cold TE buffer and membranes permeabilized by mixing cells with 1 mL ice-cold 77% (v/v) ethanol. Suspensions were kept on ice and periodic acid (Riedel-de Haën) in ddH_2_O was added to the samples to a final concentration of 10 mg/mL.

Cells were kept on ice for at least 30 minutes, then harvested by centrifugation and re-suspended in 1 mL PBS-E following two wash steps with the same buffer. Suspensions were kept on ice prior to the dropwise addition of 25 µL Schiff’s reagent (Sigma) heated to 90°C. Following three inversions, samples were incubated at room temperature with slight agitation (150 RPM) for 30 minutes. Excess dye was removed by washing the cells three times with PBS-E and bacterial pellets were finally re-suspended in GTE and stored at 4°C. Imaging occurred within 48 hours of staining.

### Immunofluorescent labelling of FtsZ

Z-rings were labelled using an established protocol, with minor modifications^68^. Briefly, 15 mL exponentially growing cells (OD_450 ∼_ 0.1 – 0.2) were fixed directly^66^ by placing the cells in tubes containing 400 µL 1 M NaPO_4_ pH 7.4, and 3 mL ice-cold fixative (16% (v/v) formaldehyde, 0.06% (v/v) glutaraldehyde in PBS). Fixation was carried out at room temperature for 15 minutes with agitation (100 RPM), after which samples were placed on ice for another 30 minutes. Fixed cells were harvested by centrifugation (4 500 x g, 5 mins, 4°C), washed three times with PBS-E and stored in the same buffer at 4°C. Immunolabelling procedures were carried out within 48 hours of fixation. All subsequent steps were performed in 150 µL volumes and centrifugation steps were at 4 500 x g for 5 mins at 4°C.

Permeabilization was achieved by re-suspending fixed cells in 150 µL PBS containing 0.1% (v/v) Triton X-100 (PBS-T). Bacterial cells were incubated at room temperature for 45 minutes, washed three times with PBS-E and re-suspended in PBS containing 100 µg/mL lysozyme and 5 mM EDTA. Samples were incubated for 45 minutes at room temperature before cells were collected and washed three times in PBS-E.

Cells were incubated in blocking buffer (2% BSA (w/v) in PBS) at 37°C for 30 minutes. They were collected by centrifugation and FtsZ was labelled using α-FtsZ serum (Agrisera) diluted in blocking buffer (1:500). Incubation in primary antibody occurred at 37°C for 1.5 hours with slight agitation (200 RPM). Cells were collected and washed three times in PBS-T, and then incubated in goat anti-rabbit IgG conjugated to Alexafluor 488 fluorophore (Invitrogen) diluted 1:200 in blocking buffer. Samples were incubated at 37°C for 30 minutes with slight agitation (200 RPM), after which cells were washed three times in PBS-T, washed once in PBS and finally re-suspended in 100 µL GTE. Bacteria were kept at 4°C and were imaged within 24 hours of labelling FtsZ.

### Sample mounting

Approximately 20 µL of cells were air-dried either on agarose pads (1% (w/v) in PBS) or on slides coated with 0.1% (w/v) poly-l-lysine solution (Sigma) and bacterial nucleoids stained by topping cells with a drop of Fluoroshield™ containing 1.5 µg/mL 4’,6-diamidino-2-phenylindole (DAPI, Sigma). Cover slips were placed over the samples and kept at room temperature for ∼ 30 minutes before cells were imaged.

### Confocal image acquisition

Cells were visualized using a Zeiss LSM780 ElyraPS1 confocal microscope equipped with a 100X alpha Plan-Apochromat (N.A. = 1.46) M27 objective (Carl Zeiss). For nucleoid, membrane and viability staining the 405 nm, 488 nm and 561 nm laser lines were used to excite DAPI, FM 4-64FX and the fixable LIVE/DEAD dye, respectively. Glycogen bodies were examined by exciting PAS-stained polysaccharides with a 488 nm laser, which was also used for the excitation of Alexa 488 fluorophores used for Z-ring visualizations.

Emission was detected with a GaAsP detector in the spectral ranges of 415 nm to 498 nm (λ_em_ = 450 nm) for DAPI, 572 nm to 733 nm (λ_em_ = 650 nm) for FM 4-64FX, 498 nm to 555 nm (λ_em_ = 530 nm) for the LIVE/DEAD dye, 570 nm to 695 nm (λ_em_ = 630 nm) for PAS-stained polyglucans and 499 nm to 570 nm (λ_em_ = 535 nm) for Alexa 488-conjugated secondary antibodies. ZEN 2 (Black edition, Zeiss) was used for image capturing, analysis and manipulation.

### Determination of viable and achromosomal cells

The frequencies of viable and achromosomal cells were manually determined using multichannel confocal images which were contrast-corrected and superimposed in ZEN 2. Cells completely devoid of DAPI-associated fluorescence were scored as achromosomal and bacteria were classified as non-viable only if signal from the vital stain was detected throughout the entire cell. Cells showing only polar fluorescent foci were considered viable. The frequencies of cells with Z-rings were manually determined in ZEN 2 from multichannel images and cells were scored positive if they displayed a solid band of fluorescence across the width of the cell, two adjacent fluorescent foci, or a single fluorescent focus, signifying a constricting septum. Live and fixed WT and triple mutant cells (at least 5 000 bacteria measured over a total of three biological repeats) were measured during exponential growth to establish if fixation adversely affected length measurements.

### Image processing for cell size analysis

The individual dimensions of membrane-stained bacteria were determined from confocal pictures. We devised a custom Python script (coliseg) that integrates several open-source Python packages (OpenCV^69^ bindings, NumPy^70^, Pandas^71^ and Matplotlib^72^ with seaborn^73^) to permit excellent delineation of cell borders and determination of size, being particularly effective in the analysis of adherent and filamentous cells.

### Image pre-processing

In order to convert 1144×1144×3 RGB images to binary ones containing bacterial outlines, they were first converted into 8-bit grey-scale images. Image contrasts were stretched using histogram equalisation and were then convolved with a 3×3 circular Gaussian kernel to decrease the rate of false positive contour detection. We then binarized the images with an intensity threshold T = 8; the contours were set with pixel intensity = 255, the background = 0. The images were convolved with an adaptive threshold in which the threshold value was derived from a weighted sum of the block size (block size = 15) region. Contours were subsequently dilated and then eroded to remove spurious blobs using the same kernel as for the initial Gaussian blurring. Images were then thinned such that contours are reduced to a 1-pixel wide thickness followed by a dilation. The resulting images contain the segmented bacteria.

### Contour extraction

In order to extract the bacterial outline from the binary images into a table of lists representing raw contours, we used the function findContours using a two-level hierarchy retrieval mode without chain approximation. We opted for this particular retrieval mode as it better separated clumped/dividing bacteria; the outer contour is usually the clump itself whereas the inner contours are the actual bacteria.

Since the inner contours are extracted without distinction between agglutinated/isolated bacteria, we can reasonably assume that it does not create a bias. The potential holes were then discriminated on the base of their abnormal shape and surface area. We kept only contours with a ratio convex hull length / contour length greater than .9 and a contour area greater than 110 pixels. We finally modelled the bacteria shape by fitting the contours into rectangles. The length, the width of the fitted rectangle as well as the area of the raw contour were saved into a set of tuples (length, width, area).

### Length analysis of whole populations

Since metric parameters of a cell are positively defined, we applied a log_10_-transformation to the width, length and area. To reduce the complexity, we binned the length of the bacteria with cuts at 1, 2, 4 and 32 µm. Using the counts in each bin, we compared the relative frequency of each bin. We visualised the proportion of long cell in each genotype at different time points using the binned cuts {0, 1, 2, 4, 32}. Finally, we plotted the difference of the bins for each genotype against the WT to better visualise the enrichment in each bin.

### Analysis of the fraction of elongated cells

We considered the probability of observing elongated cells (length > 4 µm) and investigated how this probability varies between genotypes. The average cell length is not sensitive to the presence of only a few elongated cells. Instead, we sought to evaluate the probability of observing one elongated cell. Thus, at each time point, the multi-level binomial model was used.

~~~
$$y_i \sim Binomial(N_i, p_i)$$
$$logit(p_i) = \alpha_i$$
$$\alpha_i \sim Normal(\mu, \sigma)$$
$$\mu \sim Normal(−2, 1)$$
$$\sigma \sim half-Normal(0, 1)$$
~~~

Where $$y_i$$ is the number of observed elongated cells, $$p_i$$ is the successful probability of observing an elongated cell, $$N_i$$ is the total number of observed cells, and $$i$$ denotes the genotype. For computational reasons, we used the $$logit$$ link function. Therefore, the intercepts $$\alpha_i$$ can be modelled using a normal prior, whose mean is centred at –2 (expit(2) = .1) - a sensible value for a small probability of elongated cell.

We used Stan^74^ and its Python interface PyStan^75^ to draw samples from the posterior joint distribution. We ran 4 independent Markov chains during 4 000 iterations (see detailed model summaries in Supplementary Table 2). To compare the proportions of elongated cells, we computed the difference of proportion between each pair of genotypes. To figure out the expected proportions, one needs to use the inverse logit function also called expit function.

In Stan, the mathematical model turns into

~~~
data {
 int<lower=0> N;        // Number of observations
 int<lower=0> G;        // Index encoding the genotype
 int n[N];                // Number of observations in genotype
 int y[N];                // Number of elongated cells in genotype
 int<lower=1, upper=G> genotype[N]; // Encoding for the genotype
}
 parameters {
  real mu;
   real<lower=0> sigma;
   vector[G] alpha;
}
 model {
  mu ∼ normal(−2, 1);
  sigma ∼ std_normal();
  alpha ∼ normal(mu, sigma);
  y ∼ binomial_logit(n, alpha[genotype]); // Likelihood in unconstrained space
}
 generated quantities {
   real<lower=0,upper=1> p[G];
   for (i in 1:G)
       p[i] = inv_logit(alpha[i]);
}
~~~

### Statistical analyses

A minimum of three independent replicates were performed for all experiments, unless stated otherwise. Statistical tests were performed using Prism 6 (GraphPad Software) or Statistica 13 (Dell Software). Normality of data was assessed by Shapiro-Wilk tests and homoscedasticity demonstrated using Brown-Forsythe variance tests. Significant differences (α = 0.05) between group means were determined by performing a one-way ANOVA with Tukey’s post hoc test. A log_10_-transformation was applied to culturability data prior to performing an analysis of variance to correct for heteroscedasticity and skewness of data.

## FUNDING

This work was supported by the Swiss/South African Joint Research Program [grant number 87391] and the National Research Foundation’s South African Research Chairs Initiative [Genetic tailoring of biopolymers] as well as by an Innovation Master’s Scholarship [UID 114278] to FvdW from the South African National Research Foundation and Department of Science and Technology.

## ACKNOWLEDGEMENTS

The authors would like to thank Professor Kirsten Skarstad from the Oslo University Hospital for the kind gift of the DnaA antibody. We would also like to express our gratitude towards Mr Michael Rosmarin for imaging some of the bacterial plates used for viable titre determinations and Mr Gert Grobler for general advice relating to data analytics.

## AUTHOR CONTRIBUTIONS

JRL conceived the project and FEvdW conceptualized parts thereof. LB created coliseg and performed cell sizing analyses. LS generated mutant strains and FEvdW performed all experimental procedures. LE assisted with experiments relating to confocal microscopy and RCMA with those pertaining to flow cytometry. JdS and FEvdW manually interpreted confocal micrographs. JRL and FEvdW wrote the manuscript. JRL and JK provided funding. GMG and SCZ provided critical feedback of the original paper. All authors were involved in reviewing and editing the final manuscript.

### Conflict of interest

None declared

## Addendum A

### Figure and table legends

**Table S1.** Mean cellular widths of the bacterial strains. Cell dimensions were determined at the indicated growth phases from confocal micrographs across three independent repeats. Values reported are mean cell widths (µm) of ≥ 550 cells examined for each experiment ± standard deviation.

|  | Mean cell width (μm) |  |  |
| --- | --- | --- | --- |
| Strain | Early exponential | Onset stationary | Late stationary |
| Wild type | 0.90 ± 0.14 | 0.78 ± 0.10 | 0.80 ± 0.12 |
| *ΔglgP* | 0.88 ± 0.15 | 0.78 ± 0.08 | 0.86 ± 0.12 |
| *ΔglgX* | 0.87 ± 0.15 | 0.77 ± 0.09 | 0.85 ± 0.13 |
| *ΔglgP/ΔglgX* | 0.89 ± 0.16 | 0.82 ± 0.12 | 0.86 ± 0.12 |
| *ΔmalP* | 0.88 ± 0.21 | 0.78 ± 0.12 | 0.83 ± 0.11 |
| *ΔmalP/ΔglgP* | 0.88 ± 0.20 | 0.81 ± 0.14 | 0.83 ± 0.11 |
| *ΔmalP/ΔglgX* | 0.93 ± 0.25 | 0.80 ± 0.12 | 0.94 ± 0.14 |
| *ΔmalP/ΔglgP/ΔglgX* | 0.90 ± 0.19 | 0.83 ± 0.14 | 0.90 ± 0.12 |

**Table S2.** Summary of the draws from the posterior predictive distribution. The indices [1 to 8] refer to the respective genotypes (‘*ΔglgP*’, ‘*ΔglgP/ΔglgX*’, ‘*ΔglgX*’, ‘*ΔmalP*’, ‘*ΔmalP/ΔglgP*’, ‘*ΔmalP/ΔglgP/ΔglgX*’, ‘*ΔmalP/ΔglgX*’, ‘WT’). The quantities p are the predicted proportions of elongated cells and the alphas are the intercepts for the logistic regressions. The mean and standard deviation of each parameter are reported along with their uncertainty whose quartiles are reported in the table. To measure the degree of overlap between two comparable parameters of interest, one can compare the extreme values of the parameter distribution (2.5 and 97.5 %) of the two parameters. If their extreme values do not overlap, it is reasonable to assume they are different. For further information about the sampling, see dedicated references discussing Bayesian inference using MCMC sampling.

| Early exponential phase |  |  |  |  |  |  |  |  |  |  |
| --- | --- | --- | --- | --- | --- | --- | --- | --- | --- | --- |
| Inference for Stan model: anon_model_726b2a34be7677656181f369bb671718. 4 chains, each with iter = 4000; warmup = 2000; thin = 1; post-warmup draws per chain = 2000, total post-warmup draws = 8000. |  |  |  |  |  |  |  |  |  |  |
|  | Mean | SE_Mean | SD | 2.5% | 25% | 50% | 75% | 97.5% | N_eff | Rhat |
| mu | -1.75 | 2.50E-03 | 0.22 | -2.2 | -1.89 | -1.75 | -1.62 | -1.3 | 7929 | 1.0 |
| sigma | 0.63 | 2.20E-03 | 0.19 | 0.37 | 0.49 | 0.59 | 0.72 | 1.12 | 7951 | 1.0 |
| alpha[1] | -2.17 | 2.70E-04 | 0.03 | -2.24 | -2.19 | -2.17 | -2.15 | -2.1 | 15971 | 1.0 |
| alpha[2] | -2.0 | 2.90E-04 | 0.03 | -2.07 | -2.02 | -2.00 | -1.98 | -1.93 | 14028 | 1.0 |
| alpha[3] | -2.15 | 2.80E-04 | 0.03 | -2.21 | -2.17 | -2.15 | -2.13 | -2.08 | 14579 | 1.0 |
| alpha[4] | -1.33 | 2.80E-04 | 0.03 | -1.4 | -1.35 | -1.33 | -1.31 | -1.27 | 14777 | 1.0 |
| alpha[5] | -1.2 | 2.80E-04 | 0.03 | -1.27 | -1.23 | -1.2 | -1.18 | -1.14 | 15136 | 1.0 |
| alpha[6] | -1.43 | 2.50E-04 | 0.03 | -1.49 | -1.45 | -1.43 | -1.41 | -1.37 | 14433 | 1.0 |
| alpha[7] | -1.11 | 2.40E-04 | 0.03 | -1.17 | -1.13 | -1.11 | -1.09 | -1.05 | 16349 | 1.0 |
| alpha[8] | -2.55 | 3.40E-04 | 0.04 | -2.63 | -2.58 | -2.55 | -2.52 | -2.47 | 13723 | 1.0 |
| p[1] | 0.1 | 2.50E-05 | 3.20E-03 | 0.1 | 0.1 | 0.1 | 0.1 | 0.11 | 15967 | 1.0 |
| p[2] | 0.12 | 3.00E-05 | 3.60E-03 | 0.11 | 0.12 | 0.12 | 0.12 | 0.13 | 14064 | 1.0 |
| p[3] | 0.1 | 2.60E-05 | 3.10E-03 | 0.1 | 0.1 | 0.1 | 0.11 | 0.11 | 14589 | 1.0 |
| p[4] | 0.21 | 4.60E-05 | 5.50E-03 | 0.2 | 0.21 | 0.21 | 0.21 | 0.22 | 14825 | 1.0 |
| p[5] | 0.23 | 4.90E-05 | 6.00E-03 | 0.22 | 0.23 | 0.23 | 0.23 | 0.24 | 15134 | 1.0 |
| p[6] | 0.19 | 3.80E-05 | 4.60E-03 | 0.18 | 0.19 | 0.19 | 0.2 | 0.2 | 14461 | 1.0 |
| p[7] | 0.25 | 4.50E-05 | 5.80E-03 | 0.24 | 0.24 | 0.25 | 0.25 | 0.26 | 16320 | 1.0 |
| p[8] | 0.07 | 2.30E-05 | 2.70E-03 | 0.07 | 0.07 | 0.07 | 0.07 | 0.08 | 13726 | 1.0 |
| lp__ | -2.40E+04 | 0.04 | 2.33 | -2.40E+04 | -2.40E+04 | -2.40E+04 | -2.40E+04 | -2.40E+04 | 2899 | 1.0 |

|  | Mean | SE_Mean | SD | 2.5% | 25% | 50% | 75% | 97.5% | N_eff | Rhat |
| --- | --- | --- | --- | --- | --- | --- | --- | --- | --- | --- |
| mu | -4.43 | 6.70E-03 | 0.54 | -5.34 | -4.8 | -4.48 | -4.11 | -3.22 | 6530 | 1.0 |
| sigma | 1.5 | 4.80E-03 | 0.38 | 0.91 | 1.22 | 1.44 | 1.72 | 2.39 | 6272 | 1.0 |
| alpha[1] | -7.26 | 3.80E-03 | 0.38 | -8.08 | -7.5 | -7.24 | -7 | -6.59 | 10094 | 1.0 |
| alpha[2] | -5.19 | 1.10E-03 | 0.13 | -5.44 | -5.27 | -5.19 | -5.1 | -4.95 | 12364 | 1.0 |
| alpha[3] | -6.16 | 2.00E-03 | 0.21 | -6.59 | -6.3 | -6.15 | -6.02 | -5.78 | 10211 | 1.0 |
| alpha[4] | -4.21 | 9.40E-04 | 0.09 | -4.4 | -4.28 | -4.21 | -4.15 | -4.03 | 10053 | 1.0 |
| alpha[5] | -4.02 | 5.80E-04 | 0.06 | -4.14 | -4.06 | -4.01 | -3.97 | -3.89 | 12291 | 1.0 |
| alpha[6] | -3.65 | 6.00E-04 | 0.06 | -3.77 | -3.69 | -3.64 | -3.6 | -3.53 | 10901 | 1.0 |
| alpha[7] | -3.94 | 6.90E-04 | 0.07 | -4.08 | -3.99 | -3.94 | -3.89 | -3.8 | 10401 | 1.0 |
| alpha[8] | -6.47 | 2.00E-03 | 0.22 | -6.93 | -6.61 | -6.46 | -6.32 | -6.07 | 12095 | 1.0 |
| p[1] | 7.50E-04 | 2.60E-06 | 2.70E-04 | 3.10E-04 | 5.50E-04 | 7.10E-04 | 9.10E-04 | 1.40E-03 | 11569 | 1.0 |
| p[2] | 5.60E-03 | 6.30E-06 | 7.00E-04 | 4.30E-03 | 5.10E-03 | 5.60E-03 | 6.00E-03 | 7.10E-03 | 12432 | 1.0 |
| p[3] | 2.10E-03 | 4.20E-06 | 4.30E-04 | 1.40E-03 | 1.80E-03 | 2.10E-03 | 2.40E-03 | 3.10E-03 | 10608 | 1.0 |
| p[4] | 0.01 | 1.40E-05 | 1.40E-03 | 0.01 | 0.01 | 0.01 | 0.02 | 0.02 | 9972 | 1.0 |
| p[5] | 0.02 | 1.00E-05 | 1.10E-03 | 0.02 | 0.02 | 0.02 | 0.02 | 0.02 | 12313 | 1.0 |
| p[6] | 0.03 | 1.50E-05 | 1.50E-03 | 0.02 | 0.02 | 0.03 | 0.03 | 0.03 | 11009 | 1.0 |
| p[7] | 0.02 | 1.30E-05 | 1.30E-03 | 0.02 | 0.02 | 0.02 | 0.02 | 0.02 | 10470 | 1.0 |
| p[8] | 1.60E-03 | 3.10E-06 | 3.40E-04 | 9.80E-04 | 1.30E-03 | 1.60E-03 | 1.80E-03 | 2.30E-03 | 12402 | 1.0 |
| lp__ | -4807 | 0.04 | 2.22 | -4812 | -4808 | -4807 | -4805 | -4804 | 3519 | 1.0 |

|  | Mean | SE_Mean | SD | 2.5% | 25% | 50% | 75% | 97.5% | N_eff | Rhat |
| --- | --- | --- | --- | --- | --- | --- | --- | --- | --- | --- |
| mu | -4.9 | 4.60E-03 | 0.34 | -5.48 | -5.13 | -4.93 | -4.72 | -4.11 | 5573 | 1.0 |
| sigma | 0.85 | 3.80E-03 | 0.27 | 0.48 | 0.65 | 0.8 | 0.99 | 1.54 | 5170 | 1.0 |
| alpha[1] | -5.87 | 2.10E-03 | 0.19 | -6.26 | -6 | -5.87 | -5.74 | -5.51 | 8616 | 1.0 |
| alpha[2] | -5.96 | 2.30E-03 | 0.22 | -6.42 | -6.11 | -5.95 | -5.81 | -5.56 | 9347 | 1.0 |
| alpha[3] | -5.33 | 1.30E-03 | 0.13 | -5.59 | -5.42 | -5.33 | -5.24 | -5.09 | 10357 | 1.0 |
| alpha[4] | -5.39 | 1.70E-03 | 0.17 | -5.74 | -5.5 | -5.38 | -5.28 | -5.07 | 10335 | 1.0 |
| alpha[5] | -4.87 | 1.20E-03 | 0.13 | -5.14 | -4.96 | -4.87 | -4.78 | -4.62 | 12063 | 1.0 |
| alpha[6] | -4.27 | 1.00E-03 | 0.1 | -4.46 | -4.33 | -4.27 | -4.2 | -4.08 | 8796 | 1.0 |
| alpha[7] | -4.13 | 7.70E-04 | 0.08 | -4.29 | -4.19 | -4.13 | -4.08 | -3.98 | 10507 | 1.0 |
| alpha[8] | -5.58 | 1.60E-03 | 0.17 | -5.92 | -5.69 | -5.57 | -5.47 | -5.26 | 10944 | 1.0 |
| p[1] | 2.90E-03 | 5.70E-06 | 5.40E-04 | 1.90E-03 | 2.50E-03 | 2.80E-03 | 3.20E-03 | 4.00E-03 | 8902 | 1.0 |
| p[2] | 2.60E-03 | 5.70E-06 | 5.70E-04 | 1.60E-03 | 2.20E-03 | 2.60E-03 | 3.00E-03 | 3.80E-03 | 10064 | 1.0 |
| p[3] | 4.80E-03 | 5.90E-06 | 6.10E-04 | 3.70E-03 | 4.40E-03 | 4.80E-03 | 5.20E-03 | 6.10E-03 | 10642 | 1.0 |
| p[4] | 4.60E-03 | 7.50E-06 | 7.80E-04 | 3.20E-03 | 4.10E-03 | 4.60E-03 | 5.10E-03 | 6.20E-03 | 10770 | 1.0 |
| p[5] | 7.70E-03 | 9.10E-06 | 1.00E-03 | 5.80E-03 | 7.00E-03 | 7.60E-03 | 8.30E-03 | 9.80E-03 | 12261 | 1.0 |
| p[6] | 0.01 | 1.40E-05 | 1.30E-03 | 0.01 | 0.01 | 0.01 | 0.01 | 0.02 | 8950 | 1.0 |
| p[7] | 0.02 | 1.20E-05 | 1.20E-03 | 0.01 | 0.01 | 0.02 | 0.02 | 0.02 | 10450 | 1.0 |
| p[8] | 3.80E-03 | 5.90E-06 | 6.30E-04 | 2.70E-03 | 3.40E-03 | 3.80E-03 | 4.20E-03 | 5.20E-03 | 11358 | 1.0 |
| lp__ | -2910 | 0.04 | 2.35 | -2915 | -2911 | -2910 | -2908 | -2906 | 3370 | 1.0 |

Samples were drawn using NUTS at Thu Mar 19 18:19:40 2020. For each parameter, n\_eff is a crude measure of effective sample size, and Rhat is the potential scale reduction factor on split chains (at convergence, Rhat = 1).

**Figure S1a. Representative confocal images of *E. coli* cells in early exponential phase.** Cell membranes, DNA and viability were assessed simultaneously using multi-color fluorescent staining. The cell periphery is shown in white, DNA in blue and proteins tagged by the viability dye are shown in yellow. Yellow arrows demarcate invaginating septa, with lipids in the new cell wall tagged by the membrane dye. White arrows indicate putative inclusion bodies, highlighted by the amine-reactive viability stain.

**Figure S1b. Representative confocal images of *E. coli* cells in early exponential phase.** Cell membranes, DNA and viability were assessed simultaneously using multi-color fluorescent staining. The cell periphery is shown in white, DNA in blue and proteins tagged by the viability dye are shown in yellow. Yellow arrows demarcate invaginating septa, with lipids in the new cell wall tagged by the membrane dye. White arrows indicate putative inclusion bodies, highlighted by the amine-reactive viability stain.

**Figure S2a. Representative confocal images of *E. coli* cells entering stationary phase.** The cell periphery is shown in white, DNA in blue and polypeptides in yellow. Arrows indicate foci which appear to be protein masses, illuminated upon excitation of the viability stain.

**Figure S2b. Representative confocal images of *E. coli* cells entering stationary phase.** The cell periphery is shown in white, DNA in blue and polypeptides in yellow. Arrows indicate foci which appear to be protein masses, illuminated upon excitation of the viability stain.

**Figure S3a. Representative confocal images of *E. coli* strain in late stationary phase.** The cell periphery was assessed alongside DNA and proteins, shown here in white, blue, and yellow, respectively. Arrows point to protein aggregates, highlighted by the amine-reactive dye used to assay viability.

**Figure S3b. Representative confocal images of *E. coli* strain in late stationary phase.** The cell periphery was assessed alongside DNA and proteins, shown here in white, blue, and yellow, respectively. Arrows point to protein aggregates, highlighted by the amine-reactive dye used to assay viability.

**Figure S4. Length analysis of *E. coli* strains.** (**a-c**) Weighted histograms showing cell lengths of the WT strain during (**a**) early exponential, (**b**) onset stationary and (**c**) late stationary phase. The 4 µm cut-off is indicated by a vertical dotted line. Difference plots following these images depict differences in bin counts between each mutant and the WT. 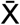
 represents mean cell lengths (µm) ± standard deviation.

**Figure S5. Immunoblot analysis of relative FtsZ levels.** Protein extracts were prepared from cells (**a**) growing exponentially, (**b**) approaching stationary phase or (**c**) cultured to late stationary phase. Ten µg total protein was loaded in each lane, separated by 10% (w/v) SDS-PAGE and blotted onto nitrocellulose membranes, before being probed with an α-FtsZ antibody. The arrow indicates the position of FtsZ (∼40 kDa) as estimated from the protein molecular size marker (PageRuler Plus, Thermo Fisher Scientific).

**Figure S6. Confocal and phase contrast imagery of PAS-stained *E. coli* cells entering stationary phase.** PAS staining procedures were performed as described and polyglucan displayed in orange, while DAPI-stained DNA is shown in blue. White arrows point to glycogen aggregates.

**Figure S7. Confocal and phase contrast imagery of PAS-stained *E. coli* cells in late stationary phase.** PAS staining procedures were performed as described and polyglucan displayed in orange, while DAPI-stained DNA is shown in blue. White arrows point to glycogen aggregates.

**Figure S8. Immunoblot examination of relative DnaA amounts.** Soluble protein was extracted from cells (**a**) growing exponentially, (**b**) entering stationary phase or (**c**) grown late into stationary phase. Ten µg total protein was separated by 10% (w/v) SDS-PAGE and blotted onto nitrocellulose membranes, before being probed with an α-DnaA antibody. The arrow points to the position of DnaA (∼53 kDa) as estimated from the protein molecular size marker (PageRuler, Thermo Fisher Scientific). An extract containing 10 µg of soluble protein from a mutant *dnaA46* strain (CM742), cultured exponentially under non-permissive conditions, was included as a negative control.

## Notes

### Competing Interest Statement

The authors have declared no competing interest.

