## Supplemental Figures 1-8 for "Disruption of glycogen metabolism alters cell size in *Escherichia coli*"

a

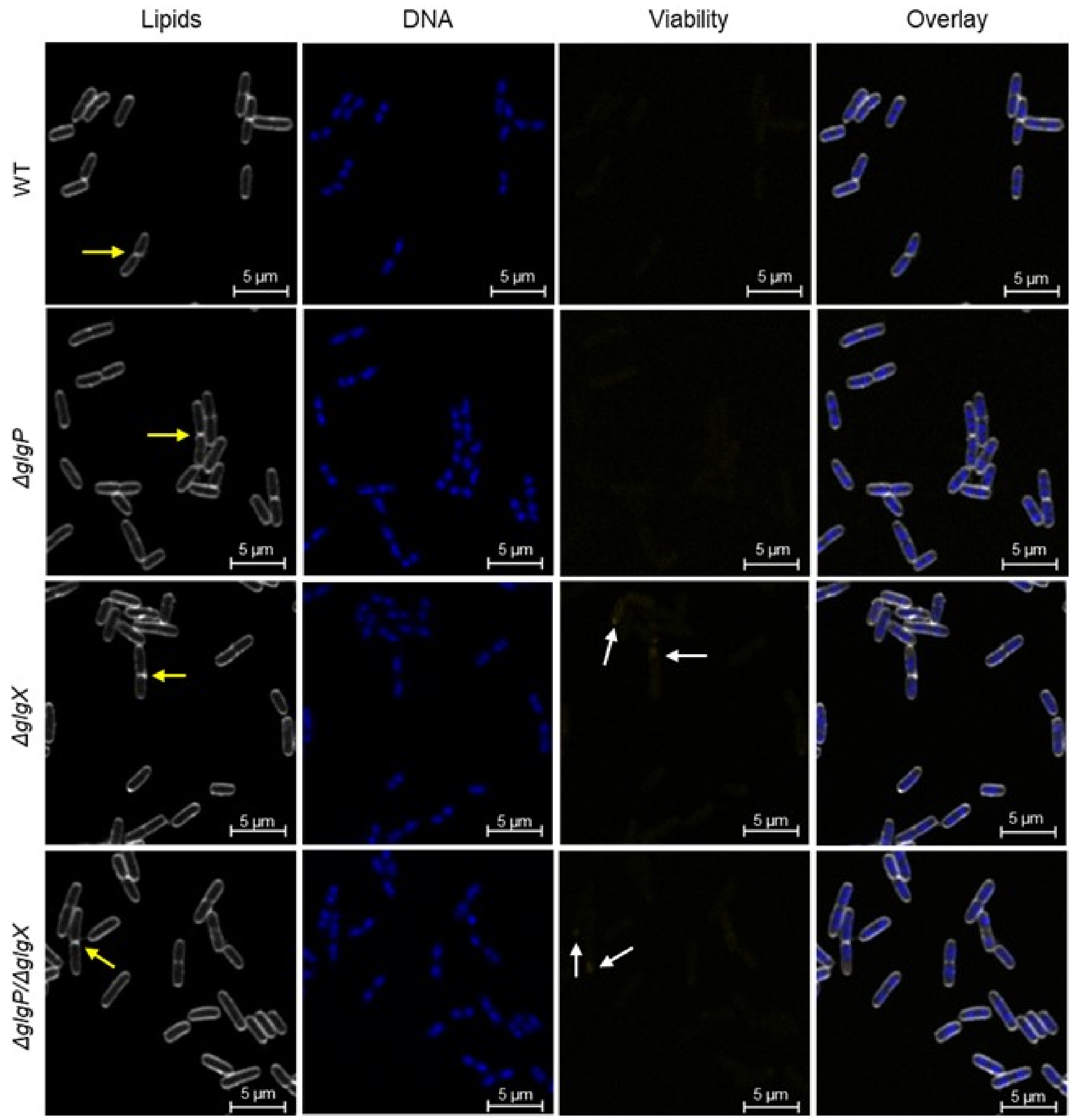


Supplemental Figure 1a

b

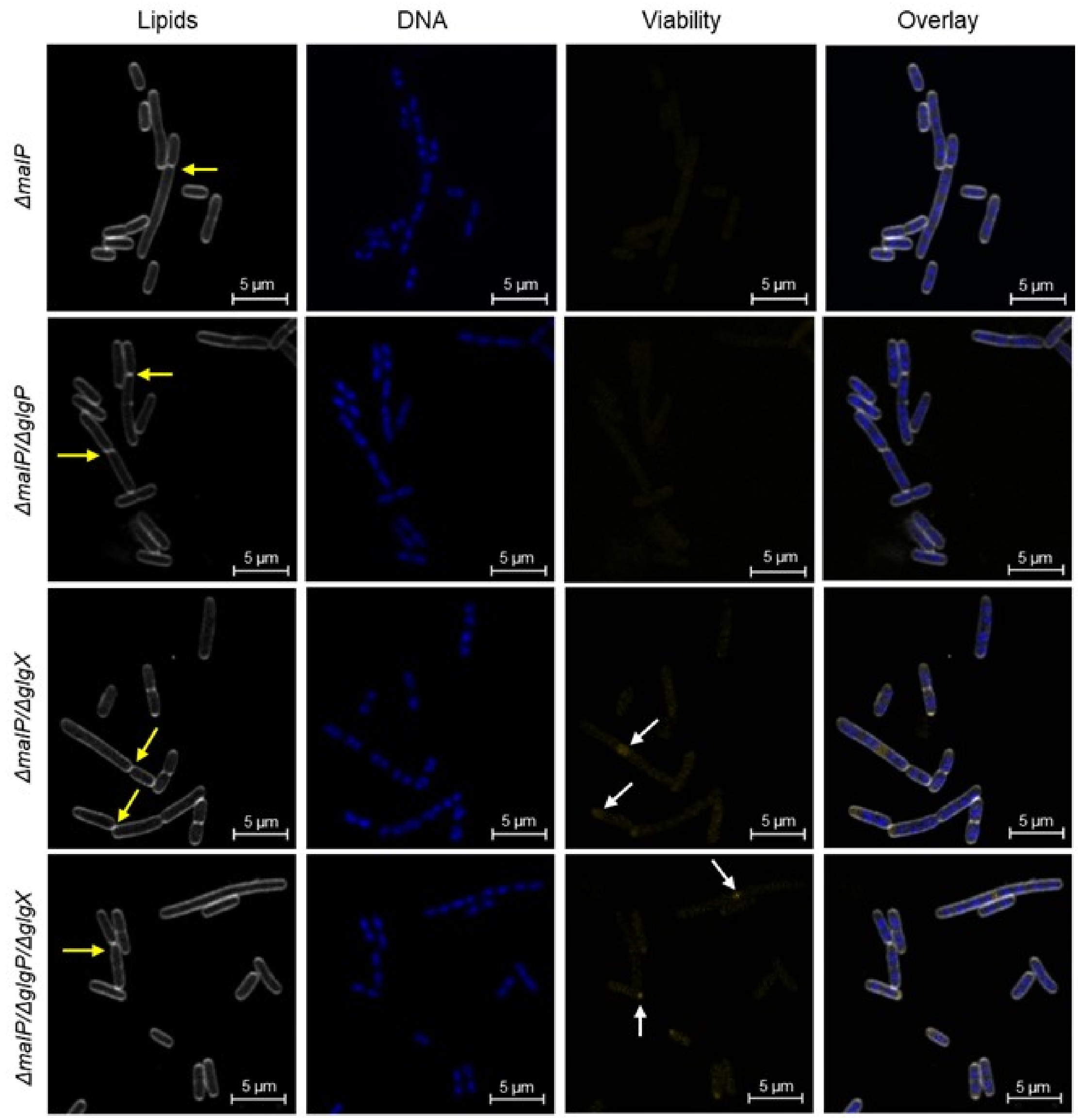


Supplemental Figure 1b

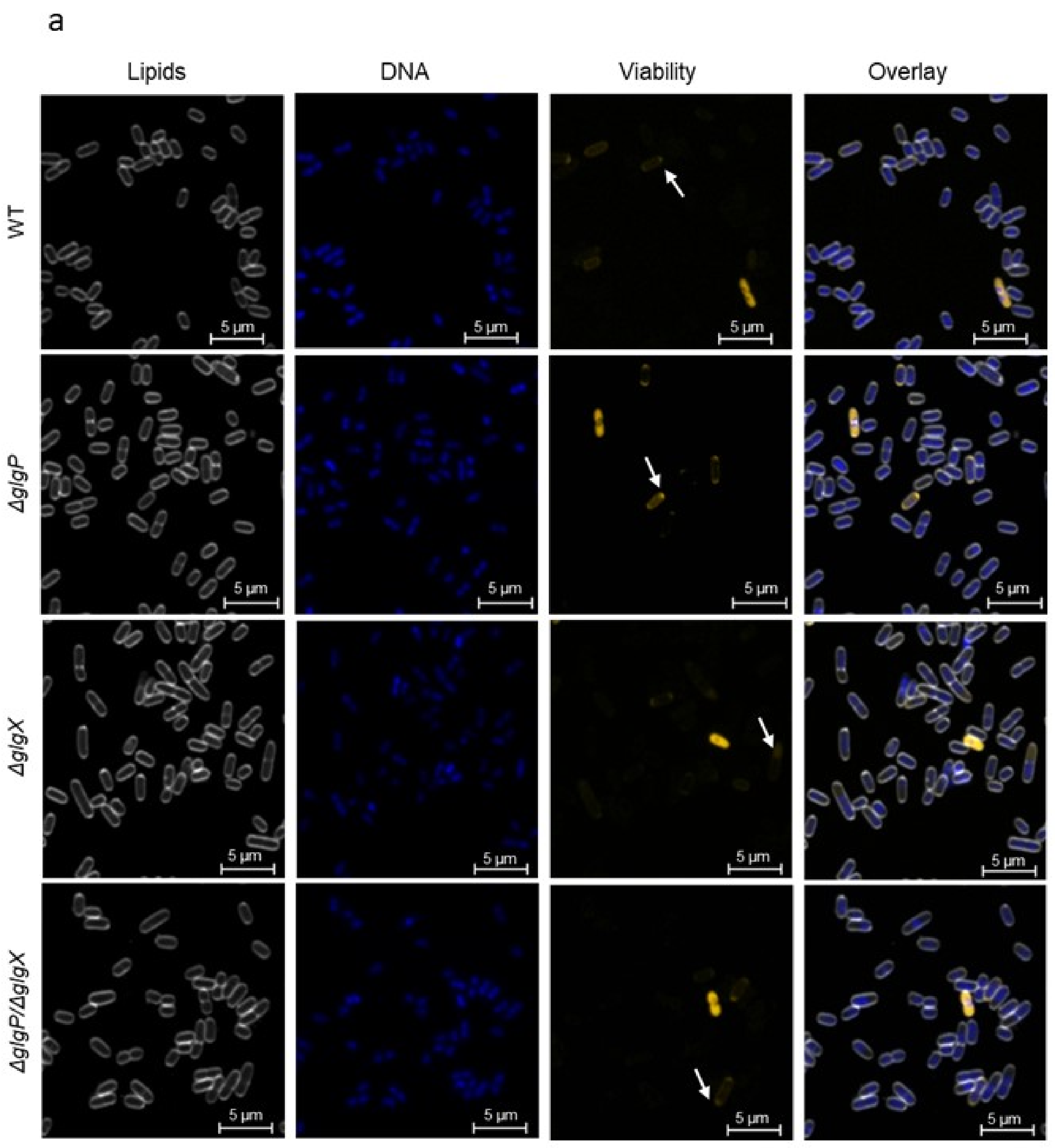


Supplemental Figure 2a

b

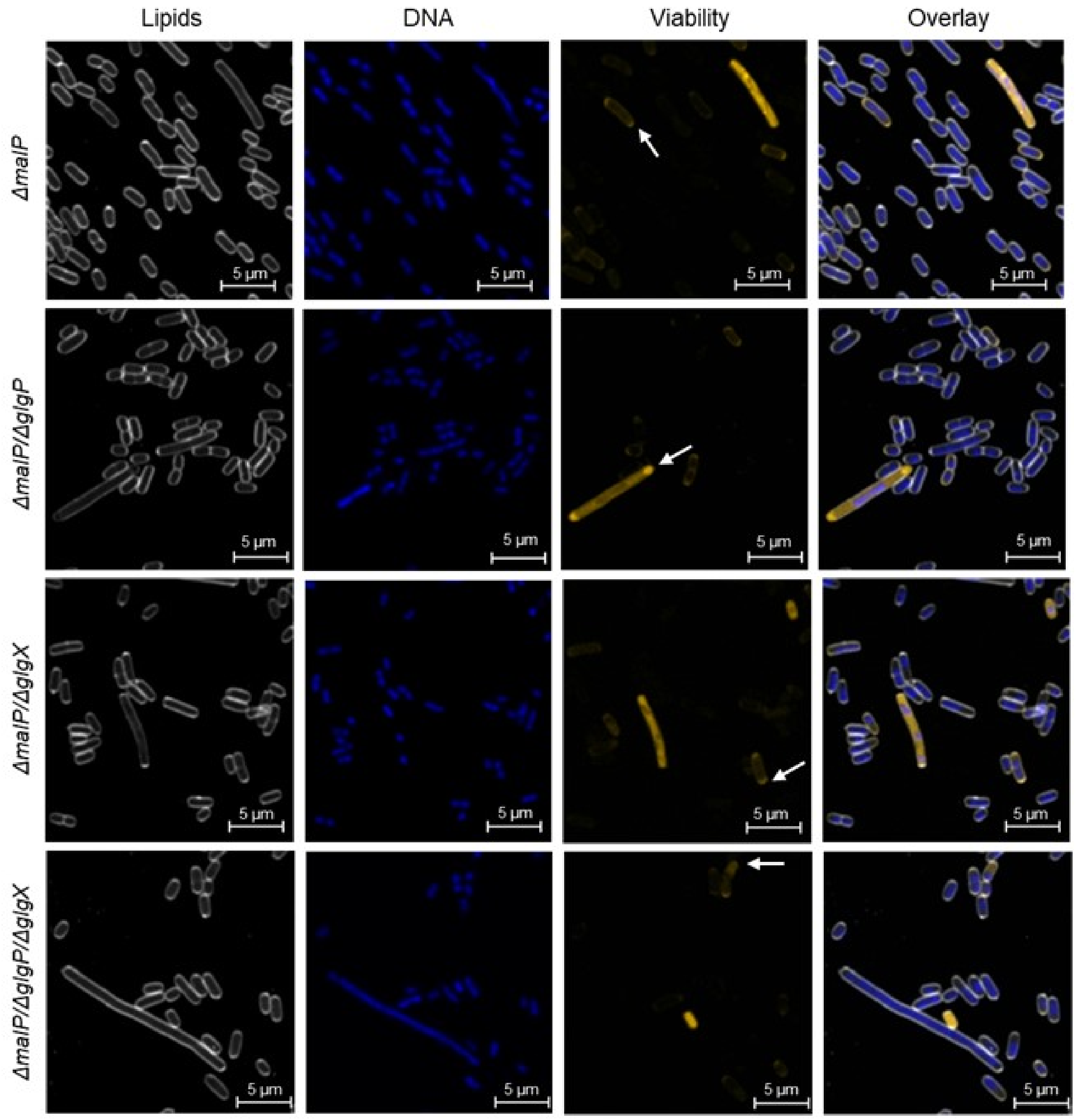


Supplemental Figure 2b

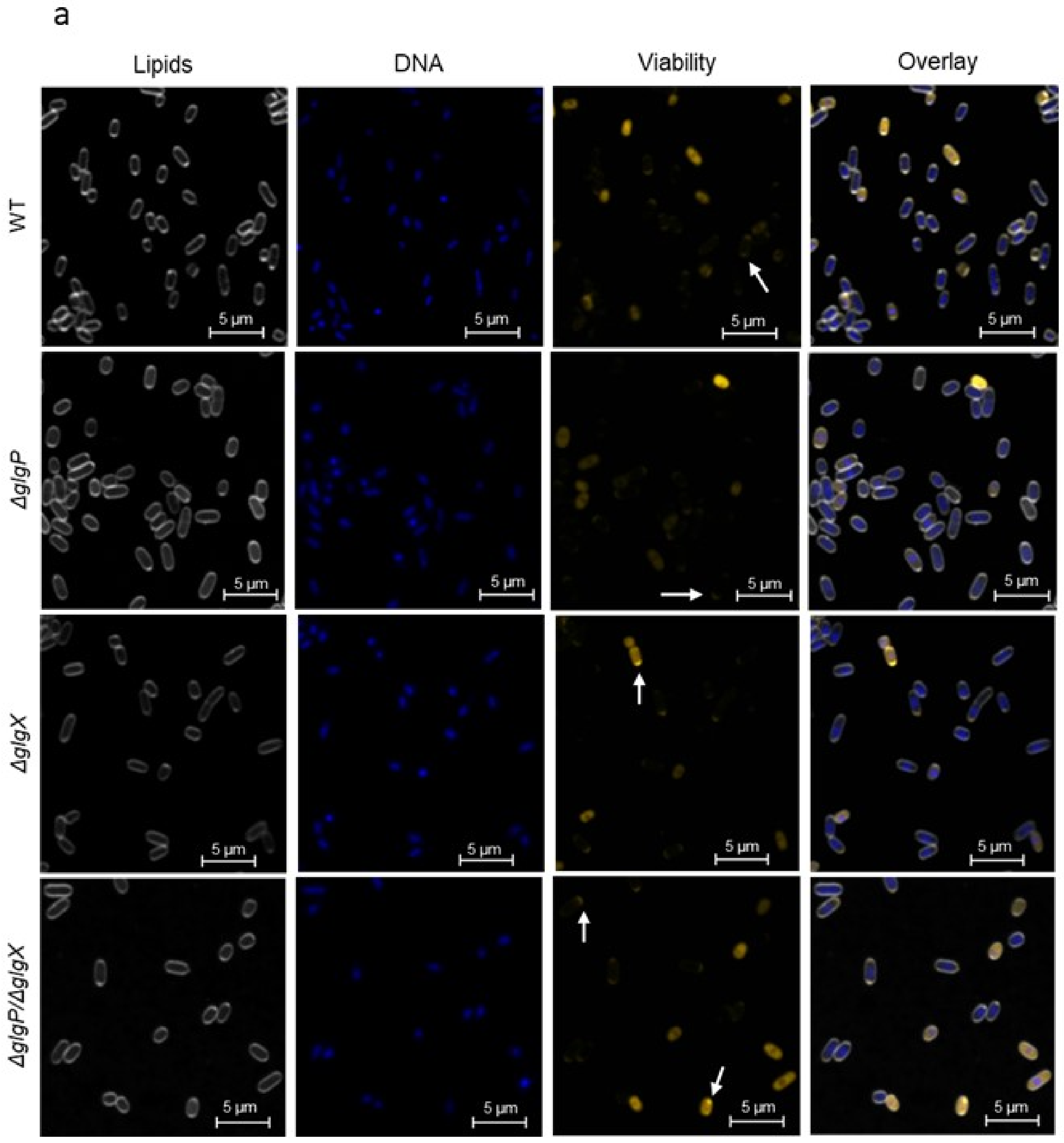


Supplemental Figure 3a

b

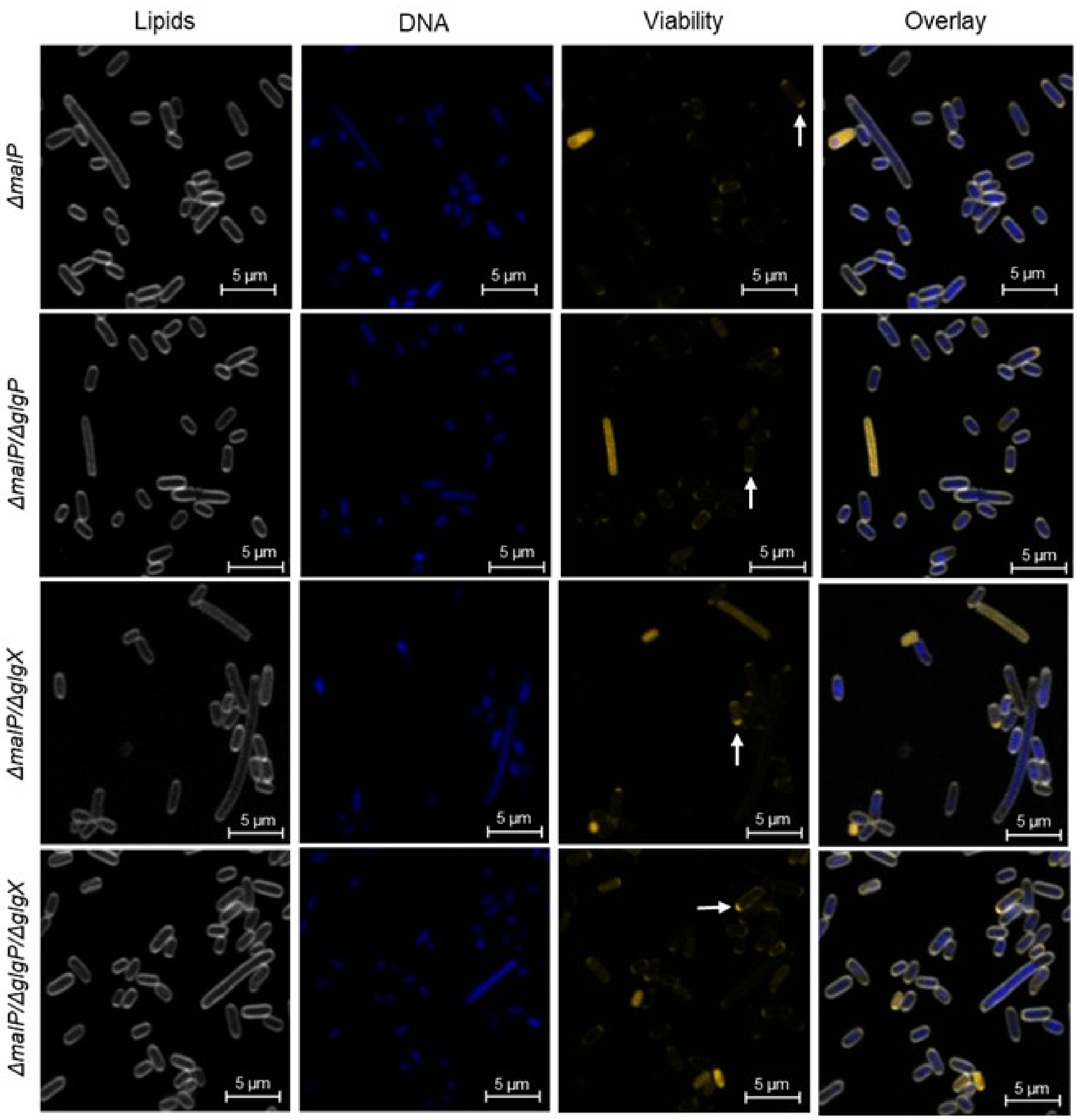


Supplemental Figure 3b

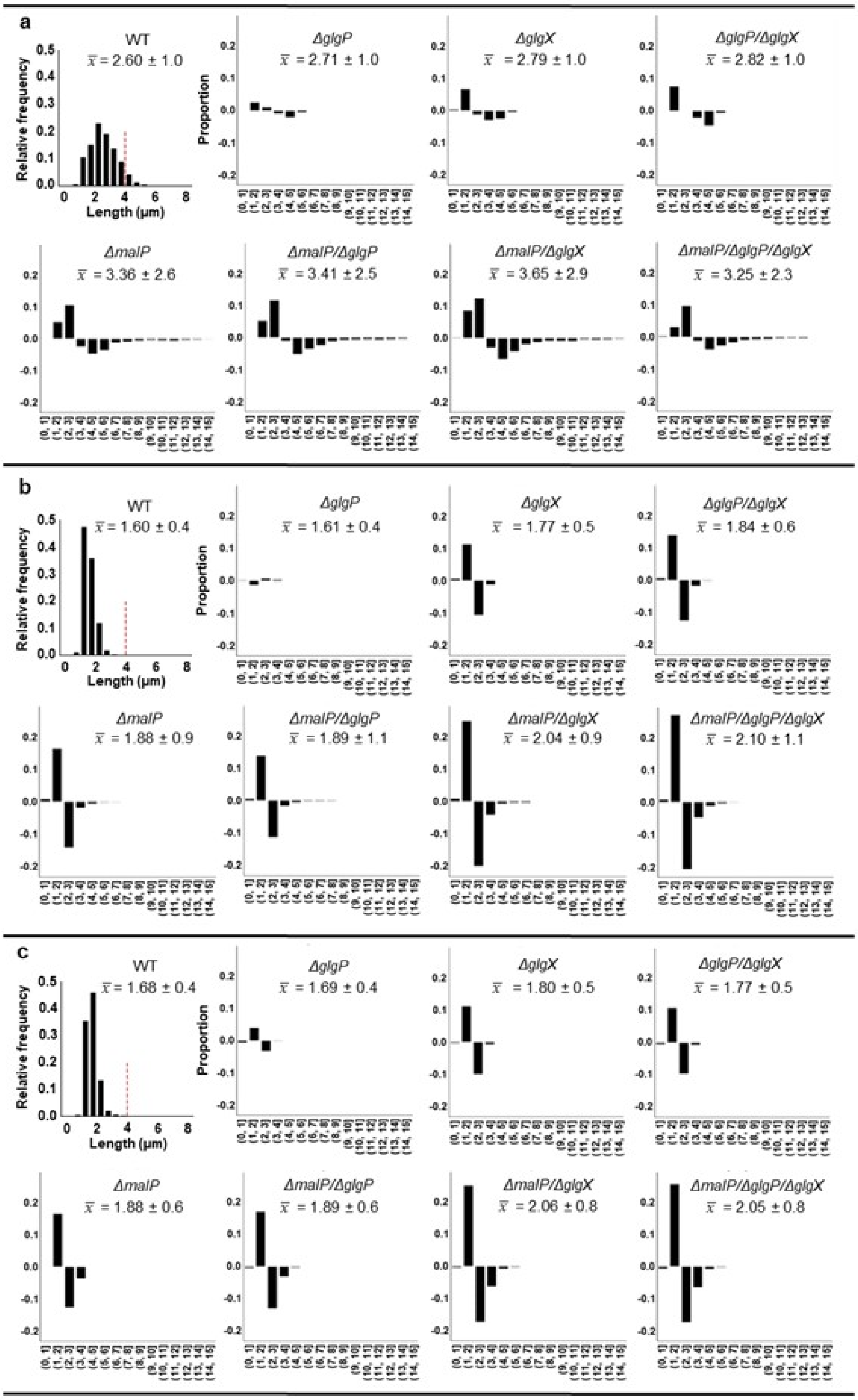


Supplemental Figure 4

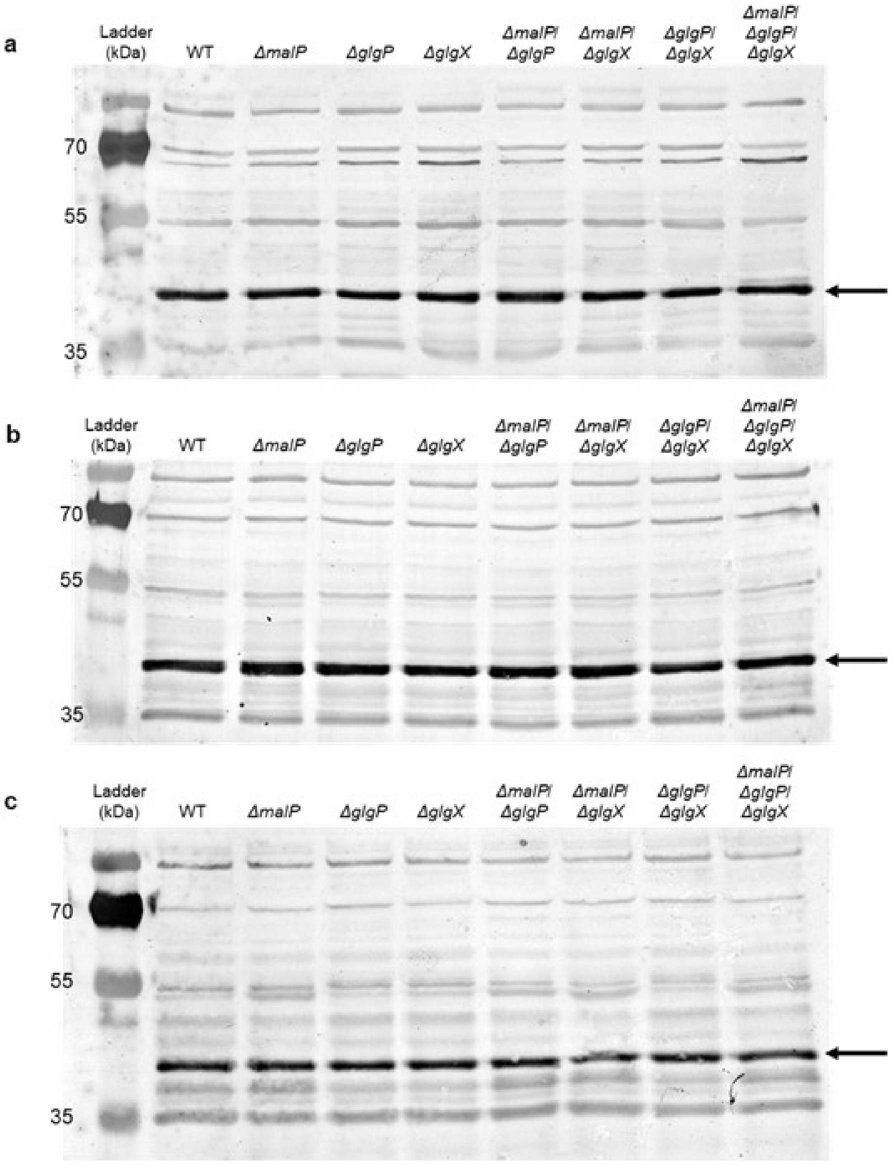


Supplemental Figure 5

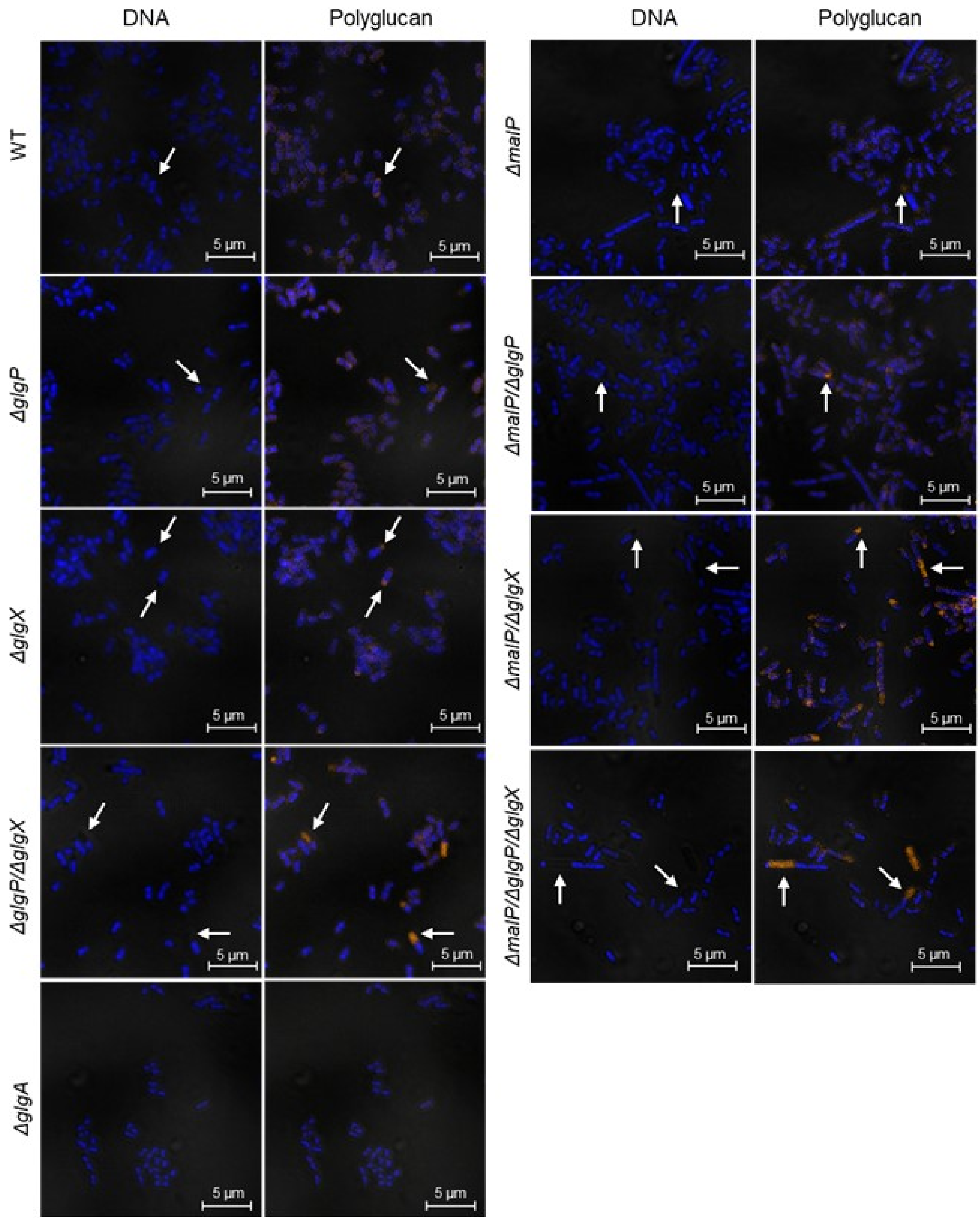


Supplemental Figure 6

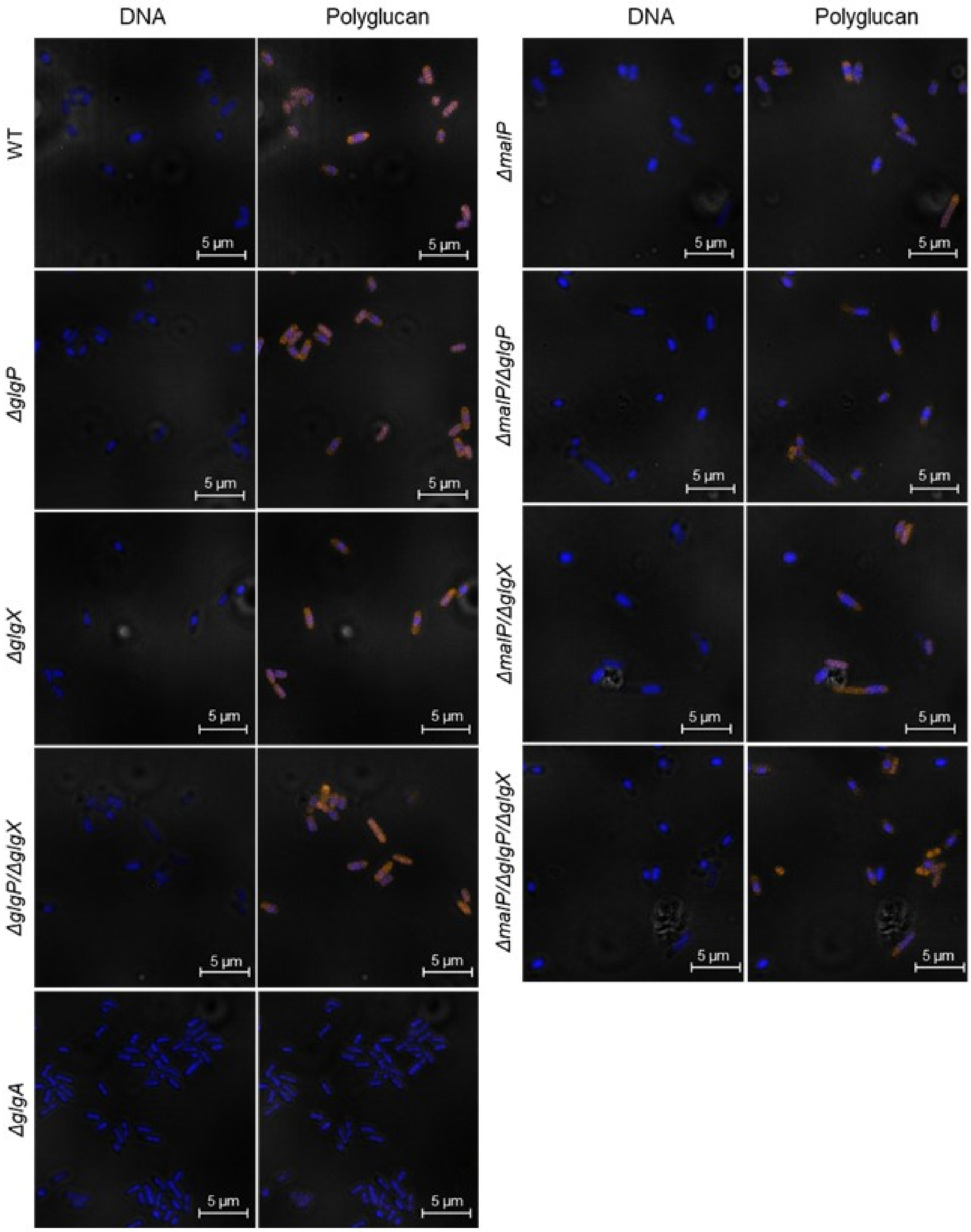


Supplemental Figure 7

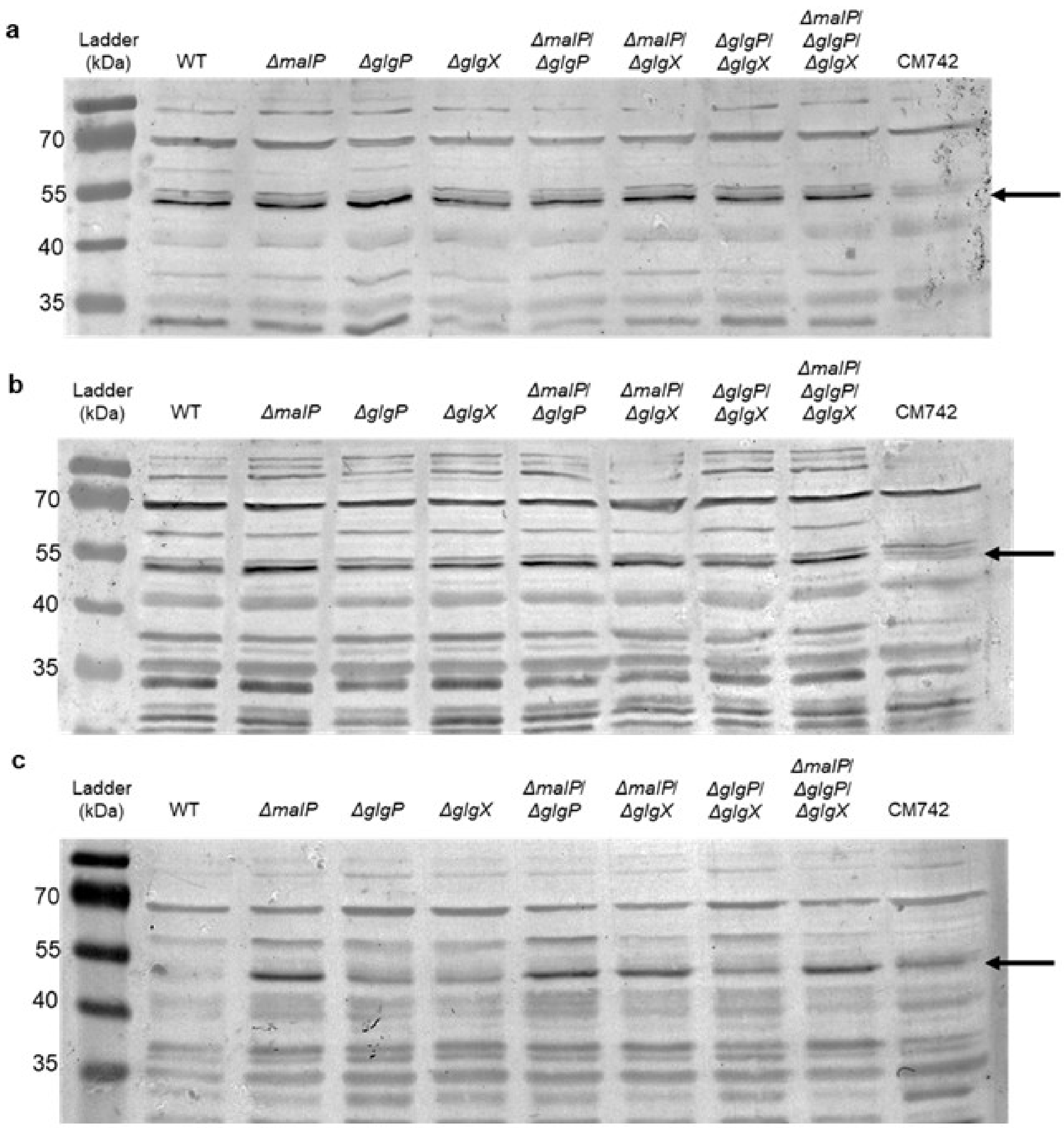


Supplemental Figure 8
